# SHERLOCK: Structured representation learning and causal inference of downstream perturbation effects

**DOI:** 10.64898/2026.09.25.754573

**Authors:** Mingxuan Zhang, Joshua D. Myers, Lingting Shi, Ross M. Giglio, Sharanya Chatterjee, José L. McFaline-Figueroa, Elham Azizi

**Affiliations:** Irving Institute for Cancer Dynamics, Columbia University, New York, NY, USA; Department of Systems Biology, Columbia University Medical Center, New York, NY, USA; Department of Molecular Pharmacology and Therapeutics, Columbia University Medical Center, New York, NY 10032, USA; Department of Biomedical Engineering, Columbia University, New York, NY, USA; Department of Computer Science, Columbia University, New York, NY, USA; Herbert Irving Comprehensive Cancer Center, Columbia University, New York, NY, USA; Data Science Institute, Columbia University, New York, NY, USA

**Author notes:** Senior and corresponding authors. Equal contribution (Mingxuan Zhang, Joshua D. Myers, Lingting Shi).

## Abstract

Understanding the effects of genetic, molecular, and experimental perturbations is essential for decoding cellular mechanisms and guiding biomedical interventions. Existing computational approaches are typically designed for individual tasks, such as predicting perturbation responses, characterizing downstream transcriptional effects, or modeling perturbation combinations, and therefore do not provide a unified framework for learning interpretable perturbation representations while enabling causal analysis of downstream effects and characterization of condition-dependent responses.

We present SHERLOCK, an interpretable deep generative framework for single-cell perturbation analysis that represents perturbation effects as structured interventions on a latent baseline cellular state. SHERLOCK learns correlated and sparse perturbation representations that organize genetic and pharmacological perturbations according to shared transcriptional responses. By formulating perturbations as interventions within a structural causal model, it enables counterfactual estimation of their downstream transcriptional effects under explicit identifiability assumptions. The same framework quantifies how perturbation responses vary across conditions and compositionally models combinatorial perturbations, enabling prediction of held-out combinations and classification of genetic interactions.

Across genome-scale CRISPR, chemical, and spatial perturbation datasets, SHERLOCK recovers perturbation relationships concordant with known biological pathways and pharmacological properties, identifies condition-dependent responses, and predicts combinatorial perturbation effects. Together, SHERLOCK provides a unified framework for interpretable and causal analysis of perturbation effects across diverse single-cell perturbation experiments.

## 1 Introduction

Large-scale single-cell perturbation technologies, including CRISPR-based screens [1, 2, 3, 4], chemical screens [5], and increasingly spatially resolved genetic perturbation screens [6, 7, 8], enable systematic characterization of cellular responses and gene function. These experiments generate high-dimensional measurements of transcriptional changes induced by diverse perturbations across heterogeneous biological and experimental conditions. The increasing scale and complexity of such datasets require computational methods that not only predict post-perturbation gene expression in and out of distribution, but importantly learn interpretable perturbation–response relationships and estimate causal effects from interventional data.

To fully realize the potential of large-scale perturbation datasets, computational methods should systematically characterize perturbation responses along several complementary dimensions: identifying perturbations that produce similar transcriptional responses and may therefore engage related biological processes; estimating their downstream transcriptional effects; determining how these effects vary across biological or experimental conditions; and characterizing the combinatorial effects produced by multiple perturbations, including departures from the effects expected from their constituent perturbations. These dimensions describe related properties of the same perturbation–response landscape and motivate computational representations that can support their joint analysis.

Existing computational approaches for single-cell perturbation analysis typically address these objectives separately, focusing on perturbation-response prediction, differential expression, representation learning, or combinatorial perturbation analysis [9, 10, 11]. These approaches have substantially advanced the analysis of perturbation screens, yet most are designed to predict or characterize perturbation-associated changes and do not support causal analysis of downstream perturbation effects. The increasing use of high-capacity models, including single-cell foundation models[12, 13], has improved predictive performance and generalization for perturbed cell gene expressions. However, predictive accuracy alone does not ensure that learned representations capture interpretable perturbation relationships or support causal analysis. A particular challenge for interpretation is identifiability because high-capacity deep generative models can represent the same observed data using different parameterizations, making the biological meaning of individual latent mechanisms ambiguous without additional structural constraints.

These limitations are particularly consequential for counterfactual inference, which asks how the same underlying cellular state would have responded under an alternative perturbation. Unlike prediction of post-perturbation expression, counterfactual estimation requires a model that separates baseline cellular variation from intervention-specific effects and specifies how alternative interventions act on the same baseline state [14, 15].

We therefore focus primarily on variational autoencoders (VAEs) because their explicit generative and posterior models provide the inference and generative operations needed for counterfactual estimation, including inference of a latent baseline state from observed control cells and generation of potential outcomes after applying alternative perturbations to that state. This generative process can be represented as a structural causal model (SCM), although the base architecture alone does not guarantee causal identification. Within this class, existing methods capture distinct but incomplete components of the desired structure. scGen[10] models perturbations as global latent displacements, whereas ContrastiveVI[16] separates shared from condition-associated variation; neither assigns individual perturbations to explicit latent mechanisms. More structured approaches, including SAMS-VAE[17], SVAE[18], and SVAE+[19], use sparse mechanism shifts to associate interventions with subsets of latent variables. Although this improves modularity and makes the models more identifiable, these formulations do not jointly represent relationships among perturbations, their downstream transcriptional effects, context-dependent responses, and combinatorial interactions within a unified framework that also supports counterfactual analysis. Thus, causal interpretation requires not only a generative model of interventions, but also structural assumptions that constrain otherwise equivalent latent parameterizations.

More broadly, interpretability remains a central challenge: learned latent representations do not necessarily correspond to biologically meaningful perturbation relationships, limiting their ability to generate mechanistic hypotheses. Methods that enforce sparsity frequently do so at the expense of predictive accuracy, whereas methods optimized for prediction tend to produce entangled representations without meaningful latent geometry. As a result, the field still lacks a unified framework that can simultaneously: (1) learn structured groupings of perturbations that reflect underlying mechanistic relationships; (2) robustly link genetic perturbations to their downstream transcriptional consequences to enable mechanistic insight; (3) explicitly model condition-dependent perturbation effects; and (4) represent combinatorial perturbations in terms of their constituent interventions, enabling prediction and interpretation of non-additive effects and genetic interactions. This framework should generalize across diverse perturbation settings, including different CRISPR modalities (CRISPR interference, CRISPR activation, CRISPR knockout) and exogenous perturbations (chemicals, cytokines), as well as across single-cell and spatial data modalities, while supporting biological discovery and hypothesis generation.

Here, we introduce SHERLOCK (Single-cell Hierarchical Embedding of peRturbations and Learning Of Causal Knowledge), a variational autoencoder framework that addresses these limitations. We represent perturbations as structured interventions on a shared baseline latent cellular state, providing both a generative and a structural causal model (SCM) formulation that enables counterfactual estimation. Specifically, sparse perturbation-specific interventions constrain the latent mechanisms, providing the structural basis for causal interpretation under explicit identifiability assumptions. The model learns structured relationships among perturbations to capture similarities in their transcriptional effects, links perturbations to downstream transcriptional programs, distinguishes intervention effects from variation associated with cellular and experimental conditions, and models non-additive effects arising from combinatorial interventions. Here, we define a condition as an additional experimental or biological context under which a perturbation is applied, such as drug exposure, cytokine stimulation, co-culture, or microenvironmental state. For example, in a genetic perturbation screen performed with and without chemical exposure, SHERLOCK can learn the effect of each genetic perturbation while identifying how that effect is modified by the exogenous chemical environment. By integrating these components within a single generative formulation, SHERLOCK is designed to recover the interpretable perturbation structure, estimate causal downstream transcriptional effects of perturbations, characterize interactions among interventions, and support counterfactual analysis across diverse perturbation modalities.

Across diverse genetic, pharmacological, and imaging-based perturbation datasets, SHERLOCK recovered biologically coherent relationships among perturbations that aligned with known pathway and functional annotations. The framework further identified condition-dependent perturbation responses and their downstream transcriptional effects, and generalized to perturbation combinations withheld from training, recovering established genetic-interaction categories. Together, these results demonstrate that SHERLOCK provides a unified and generalizable framework for organizing perturbations, estimating their causal downstream effects, and characterizing how these effects vary across conditions and combinations.

## 2 Results

### 2.1 SHERLOCK disentangles baseline cellular states from perturbation-specific effects

To address the challenge of disentangling perturbation-induced transcriptional changes from intrinsic cellular heterogeneity, we developed SHERLOCK, a structured generative model based on a VAE framework that models perturbation-induced shifts from a baseline cellular state and their dependence on biological and experimental conditions **(Fig. 1a)**. SHERLOCK takes single-cell gene expression data as input together with associated condition and perturbation labels. More specifically, we define condition as an exogenous effect that modifies perturbation response, for example experimental settings such as co-culture or drug concentrations. By perturbation we refer to the molecular or CRISPR perturbations. For each perturbation, the baseline state is learned from corresponding control populations (non-targeting sgRNAs, vehicle control, unperturbed cells) from the corresponding condition, allowing perturbation effects to be defined relative to condition-matched cellular states.

**Figure 1:**
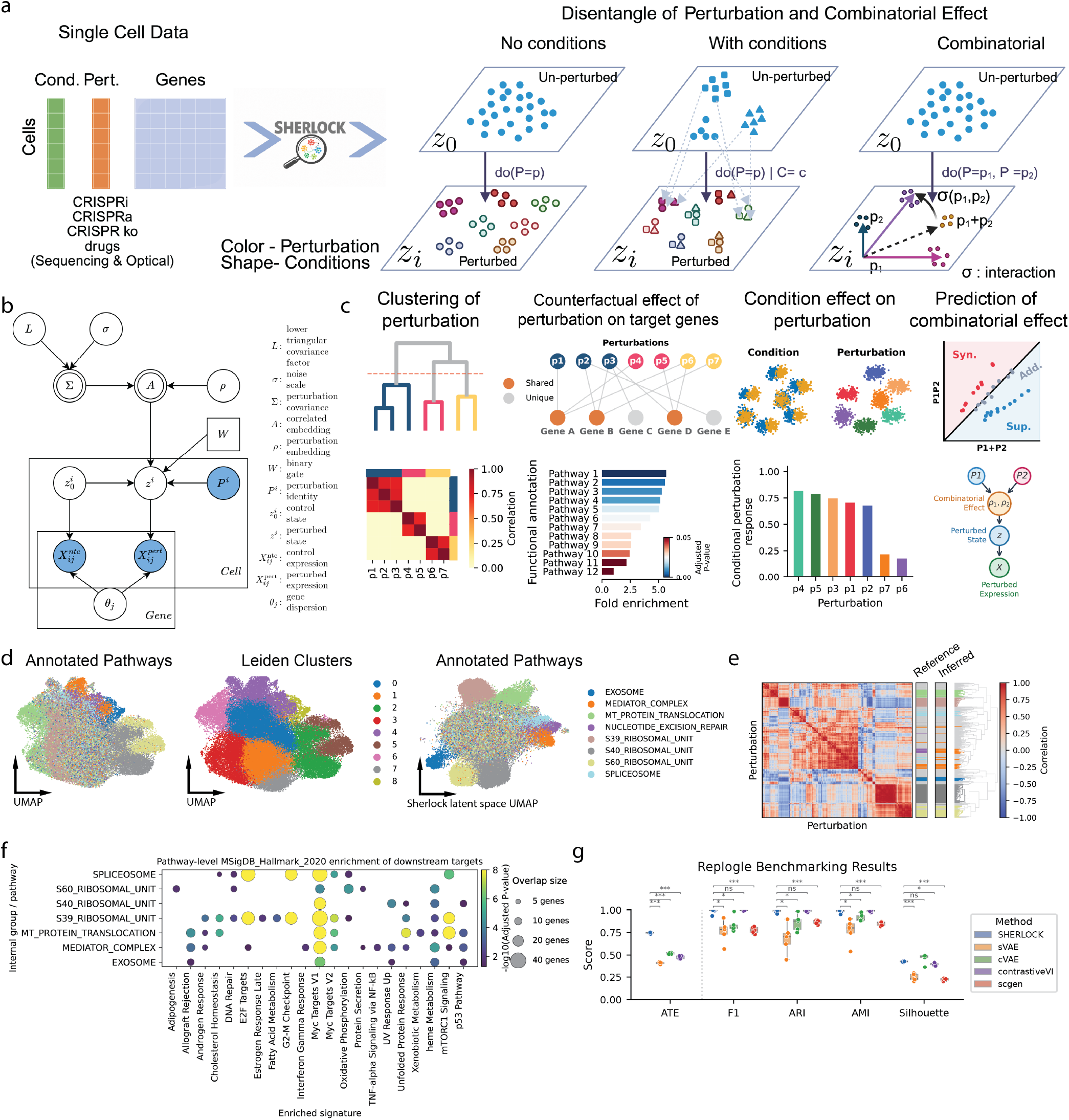
SHERLOCK: a generative framework for disentangling perturbation effects in single-cell data. (a) Overview of SHERLOCK. Single-cell gene expression data, together with perturbation and condition labels, are modeled using a structured variational autoencoder (VAE) to disentangle perturbation effects, context-dependent responses, and combinatorial interactions. (b) Plate diagram of the SHERLOCK generative process. Uncolored circles denote inferred variables, colored circles denote observed variables, and squares denote observed constants. (c) Main functionalities of SHERLOCK. (d) Left: UMAP of the original gene expression space from the Replogle dataset, colored by pathway-level annotations. Middle: UMAP of the original gene expression space, colored by Leiden clusters. Right: SHERLOCK latent representation of the same cells, colored by pathway-level annotations. (e) Correlation matrix of the learned perturbation embedding. Reference pathway annotations were annotated by Replogle et al., whereas mapped pathways were inferred from SHERLOCK groupings. (f) Gene set enrichment analysis (GSEA) of the inferred down-stream target genes for each perturbation group, highlighting pathway-relevant functional signatures. Color indicates −log_10_-transformed adjusted *p*-values, and point size represents the number of overlapping genes between the inferred target genes and enriched pathways. (g) Benchmarking of SHERLOCK on the Replogle dataset. ATE: average treatment effect; ARI: adjusted Rand index; AMI: adjusted mutual information.

SHERLOCK models unperturbed cells using a baseline latent variable (*z*_0_) that captures intrinsic transcriptional variation, and represents perturbation effects through perturbation-specific embeddings (*ρ*) that are mapped into the cellular latent space through a structured transformation **(Fig. 1a,b)**. To encode shared mechanisms across perturbations, SHERLOCK imposes a low-rank covariance structure (Σ) over the perturbation representations, allowing perturbations with related effects to be correlated in latent space. A sparse gating matrix (*W*) further restricts each perturbation to act on a subset of latent dimensions, thereby inducing modular perturbation-specific mechanisms and improving empirical identifiability of the latent representation. Together, the covariance captures relationships between perturbations, whereas the gates determine which latent dimensions each perturbation affects. The resulting perturbation-dependent latent shift is applied to the condition-matched baseline cellular state as a sparsity-regularized structural intervention (see **Methods**), producing the perturbed latent state (*z*).

This parameterization separates baseline cellular variation from perturbation-specific effects and provides a common latent representation for capturing relationships among perturbations, estimating their downstream transcriptional effects, and comparing how these effects vary across conditions. Importantly, the generative architecture also provides the operations needed for causal inference: the VAE encoder infers a latent baseline state from an observed unperturbed cell, an alternative perturbation is applied as a structured intervention on that state while holding the base state constant, and the decoder generates the corresponding potential transcriptional outcome. To improve estimation robustness, SHERLOCK constrains perturbations to act sparsely along latent dimensions and enforces covariance structures between perturbations. Under explicit assumptions on shared background noise interventions, identifiability of the noisy observation model, and common latent-coordinate alignment identifiability, the resulting counterfactual effects are identifiable (**Methods**). These conditions are theoretical assumptions rather than guarantees of finite-sample estimation; we therefore assess the empirical stability of the learned perturbation structure and counterfactual effects across independent model initializations (Fig. S1, Fig. S7). Consequently, our causal interpretations throughout the paper refer to robust approximations of the counterfactual perturbation effects (**Methods**). Across cells, SHERLOCK averages over baseline states sampled from the corresponding control population to estimate marginal perturbation effects, whereas counterfactual analysis evaluates alternative perturbations relative to a shared inferred baseline state.

Together, these components enable four complementary analyses of perturbation responses **(Fig. 1c)**. First, the learned covariance structure over perturbation embeddings enables grouping of perturbations based on shared transcriptional effects, revealing common mechanisms of action. Second, the sparse perturbation-to-latent mapping defines perturbation-specific latent shifts that can be propagated through the decoder to estimate downstream transcriptional effects. By formulating these shifts as interventions within a structural causal model, SHERLOCK further supports counterfactual estimation of perturbation effects (**Methods**). Third, SHERLOCK quantifies how perturbation responses vary across conditions, allowing systematic comparison of how perturbations reshape cellular states across biological contexts. Lastly, SHERLOCK extends to combinatorial perturbations by composing the latent shifts of their constituent perturbations with learned parent-specific scaling and a non-additive interaction residual, enabling prediction of perturbation combinations and characterization of genetic interactions relative to an additive expectation **(Fig. 1c)**.

Collectively, SHERLOCK provides a unified framework that integrates relationships among perturbations, causal estimation of their downstream transcriptional effects, condition-dependent perturbation responses, and combinatorial interactions, enabling interpretable and scalable analysis of single-cell perturbation data.

### 2.2 SHERLOCK recovers biologically coherent perturbation structure and down-stream effects in genome-scale Perturb-seq

To evaluate whether SHERLOCK can recover biologically meaningful perturbation structure from genetic perturbation data, we applied it to the genome-scale Perturb-seq dataset from Replogle et al. [20], collected with 10X Genomics Chromium technology. In a PCA-based UMAP embedding, perturbations corresponding to different biological pathways were substantially intermixed, and unsupervised Leiden clusters showed limited correspondence with the pathway-level annotations reported in the original study **(Fig. 1d)**. In contrast, the UMAP of the SHERLOCK latent space exhibits clear disentanglement of perturbation effects, with distinct groupings corresponding to annotated biological pathways.

To further assess the biological validity of these inferred groups, we examined pairwise relationships among perturbations and their downstream transcriptional consequences. The learned correlation matrix between perturbations revealed a clear block structure that was consistent with the annotated pathways **(Fig. 1e)**. SHERLOCK recovered pathway relationships broadly concordant with the pathway annotations, while grouping perturbations of the Mediator complex with nucleotide excision repair (NER) genes. Prior work indeed suggests that perturbations of transcriptional regulation and NER can produce overlapping transcriptional phenotypes, reflecting the close coupling between transcription and DNA repair [21]. Finally, gene set enrichment analysis of SHERLOCK-inferred downstream counterfactual target genes recovered pathway-relevant functional signatures for each perturbation group. This enrichment pattern is biologically coherent since MYC Targets V1, E2F Targets, and G2/M Checkpoint collectively represent a coordinated proliferative program spanning cellular growth, commitment to DNA replication, and mitotic progression, with MYC acting as an upstream regulator of many downstream transcriptional and cell-cycle pathways [22] **(Fig. 1f)**. This further supports the notion that the perturbation modules identified by SHERLOCK correspond to interpretable biological programs rather than to purely geometric separation in latent space.

We next benchmarked SHERLOCK against state-of-the-art representation learning frameworks for modeling perturbation data, including a vanilla conditional VAE, sVAE+ which uses a Beta-Bernoulli gate to learn sparse perturbation effects [19], contrastiveVI which splits representations to background and salient spaces [16], and scGen which models the perturbation effect as a linear global shift in latent space [10]. We evaluated model performance across a comprehensive suite of metrics on two primary tasks: perturbation effect estimation and latent space geometric fidelity. For perturbation-effect estimation, we compared model-derived effects with empirical marginal average treatment effects (ATEs), estimated as expression differences between cells receiving a given perturbation and the corresponding unperturbed population. We restrict this ATE comparison to generative models that explicitly learn perturbation-conditioned distributions during training. scGen is excluded from this comparison because its perturbation translation is estimated post hoc from differences between condition-specific latent means, rather than jointly learned within the generative model. For representation quality, we utilized the pathway-level annotations provided by Replogle et al. [20] as an external biological ground truth and quantified their recovery using F1 score, adjusted Rand index (ARI), adjusted mutual information (AMI), and silhouette score. SHERLOCK achieved the strongest performance across the evaluated perturbation-effect and representation metrics, indicating improved recovery of both transcriptional perturbation effects and annotated functional organization relative to the tested baselines **(Fig. 1g)**.

Finally, we evaluated robustness during retraining. Across five independently seeded runs, the learned perturbation correlation matrix was reproducible (*r* = 0.763 ± 0.022), as were the predicted counterfactual effect profiles (*r* = 0.936 ± 0.004), and within each run the predicted effect sizes tracked the measured ones (Pearson *r* = 0.891 ± 0.011). The gate co-activation pattern was less stable (*r* = 0.438 ± 0.062), as expected when latent coordinates are exchangeable between runs (**Fig. S1**). The reproducibility of factor-level functional signatures is evaluated separately in Section 2.7.

### 2.3 SHERLOCK recovers biologically coherent structure across genetic, pharmacological, and imaging-based perturbations

To assess the generalizability of SHERLOCK across modalities, we applied it to spatial single-cell genetic perturbation data generated using imaging-based RNA profiling [23] **(Fig. 2a)**. Despite the different measurement modality compared to sequencing-based assays, SHERLOCK recovered coherent perturbation groupings in its latent space **(Fig. 2b,c)**. Functional annotation using Gene Ontology (GO) analysis further supported the biological relevance of these groups; for example, group 1 was enriched for regulation of DNA-templated transcription, confirming that the inferred groupings capture shared regulatory programs **(Fig. 2d)**.

**Figure 2:**
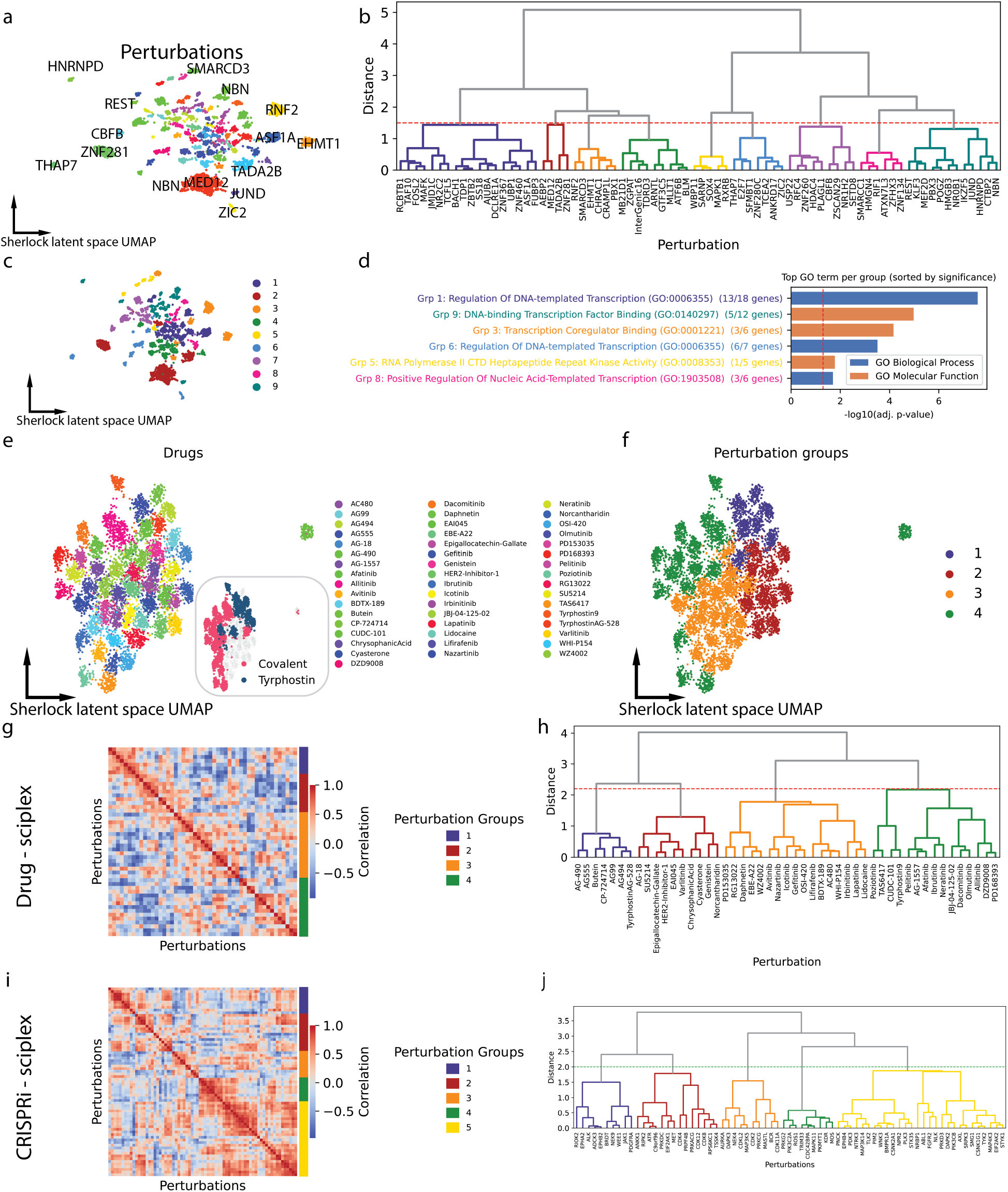
SHERLOCK identifies biologically coherent perturbation programs across sequencing- and imaging-based single-cell perturbation datasets. (a) UMAP of SHERLOCK latent space derived from spatial single-cell imaging-based RNA profiling data [23], colored by perturbation identity. (b) Hierarchical clustering of perturbations; the dashed line indicates the cutoff used to define clusters. (c) UMAP of SHERLOCK latent space colored by perturbation grouping. (d) Gene Ontology (GO) enrichment analysis of grouping of perturbations. The red dashed line indicates the significance threshold (adjusted *p <* 0.05). (e) UMAP of SHERLOCK latent space for individual drugs from the EGFR drug screen of Giglio et al. [24], colored by drug identity. (f) The same embedding colored by SHERLOCK-inferred perturbation group, revealing four major drug classes. (g) Learned correlation matrix (Σ_*p*_) of perturbation embeddings, showing block structure consistent with group-level organization. (h) Hierarchical clustering of perturbation embeddings recapitulates groupings; the red dashed line indicates the distance threshold used to define clusters. (i) Posterior correlation matrix (Σ_*p*_) of perturbation embeddings from the CRISPRi screen [25]. (j) Hierarchical clustering of CRISPRi perturbation embeddings identifies coherent groups of functionally related perturbations.

To test whether SHERLOCK generalizes across single-cell profiling techniques (10X, sci-Plex, imaging) and recovers meaningful perturbation relationships in drug-perturbation data, we applied it to the single-cell EGFR drug screen from Giglio et al. [24]. This dataset was generated using the sci-Plex framework for multiplexed single-nucleus chemical screens using single-cell combinatorial indexing (split-pool) technology [5, 26, 27], and profiled transcriptional responses to a panel of EGFR and kinase-pathway inhibitors. SHERLOCK was able to disentangle the effect of individual drugs on the latent space embedding **(Fig. 2e)** and organized the drugs into four major groups (Fig. 2f) with clearer correspondence to pharmacological properties than observed in the PCA-based representation (Fig. S2a). Notably, group 1 contained multiple tyrphostin compounds (including AG-490, AG99, AG494, Tryphostin AG-528 and AG555) (Fig. 2e, f, and h). Giglio et al. previously showed that a subset of tyrphostin-family EGFR inhibitors induce shared pro-immunogenic transcriptional program associated with increased antigen-presentation machinery and enhanced T-cell-mediated tumor killing [24].

Other groups captured distinct pharmacological properties; for example, group 4 was enriched for second-generation covalent inhibitors (including afatinib, neratinib, dacomitinib, and pelitinib), whereas group 2 contained compounds with comparatively weak transcriptional responses in the original study [24]. To further interpret the inferred perturbation structure, we annotated drugs using orthogonal chemical and pharmacological features from the original study [24] **(Fig. 2e; Fig. S2b)**. The SHERLOCK groupings showed strong concordance with known drug properties, with group 1 significantly enriched for tyrphostin-class inhibitors and group 4 significantly enriched for covalent inhibitors **(Fig. S2c)**. These results show that the learned transcriptional perturbation structure is concordant with independent chemical and pharmacological annotations.

We next applied SHERLOCK to the sci-Plex CRISPRi/a kinase perturbation screens from Shi et al. [25], which were designed to identify tumor-intrinsic regulators of cancer cell susceptibility to cytotoxic T cell-mediated killing. In this screen, we previously identified *PDGFRA* and *EPHA2* as tumor-intrinsic regulators whose perturbation altered tumor cell susceptibility to T cell-mediated killing. The CRISPRi and CRISPRa screens were modeled separately, allowing us to ask whether SHERLOCK recovered reproducible relationships among kinase perturbations within each modality. Across five independent model runs per screen, the inferred covariance structure and hierarchical clustering identified recurrent kinase groupings including *PDGFRA* and *EPHA2* in both CRISPRi and CRISPRa data **(Fig. 2i,j; S3a-c)**. Several kinases grouped with *PDGFRA* and *EPHA2* have established roles in tumor–immune interactions. These included *AXL*, which promotes resistance to cytotoxic T lymphocyte (CTL)-mediated killing in non-small cell lung cancer[28]; and *JAK1*, a central mediator of IFN-*γ* signaling whose loss impairs antigen presentation and tumor sensitivity to immune-mediated killing in melanoma [29]. Other recovered kinases have broader established roles in antitumor immunity, including *RIPK2* whose inhibition has been shown to enhance anti-PD-1 responses in pancreatic ductal adenocarcinoma [30], and *ATR*, whose inhibition promotes antitumor immune responses through DNA-damage-associated inflammatory signaling [31]. These observations are consistent with SHERLOCK grouping perturbations according to transcriptional programs relevant to tumor immune regulation. The Shi et al. study [25] directly validates PDGFRA and EPHA2 inhibition as enhancing T-cell-mediated GBM killing, while also identifying broader kinase-dependent immune-adaptation programs.

Extending beyond kinases with established links to tumor immunity, SHERLOCK also identified several kinases with less well-characterized potential roles in these programs. *PIK3CB* is a component of the PI3K signaling pathway, which has been implicated in tumor immune evasion and modulation of anti-tumor immune responses, although its direct role in regulating tumor susceptibility to T cell-mediated killing remains unclear [32]. *MAP4K3* has not been directly implicated in tumor susceptibility to T cell-mediated killing, but its established roles in amino acid sensing, autophagy, and mTOR signaling provide plausible links to cellular programs that can modulate tumor–immune interactions [33, 34]. Similarly, *PIM2* has been associated with tumor immune evasion and resistance to cancer immunotherapy, although its tumor-intrinsic role in modulating susceptibility to cytotoxic T cell-mediated killing remains largely unexplored [35]. These results show that SHERLOCK recovers reproducible kinase perturbation modules containing both established immune-regulatory genes and candidates whose roles in tumor susceptibility to T-cell pressure warrant further investigation.

### 2.4 SHERLOCK reveals context-dependent drug responses and their down-stream gene programs

Next, we sought to evaluate whether SHERLOCK captures context-dependent drug responses. We applied it to a drug perturbation dataset collected with sci-Plex, profiling 11 potential immune modulators with or without the presence of tumor antigen-specific cytotoxic T cells [25]. Drugs were applied to tumor cells and removed before T cell co-culture, enabling us to assess how drug-induced tumor cell states subsequently shape responses to T cell exposure. Compared to a PCA-based UMAP, the SHERLOCK latent representation more clearly organized cells according to both drug perturbation and T-cell co-culture condition, enabling comparison of perturbation responses across conditions and concentrations **(Fig. 3a,b)**. Because perturbation relationships are encoded through the learned covariance structure, we next examined whether these relationships changed with drug concentration and T-cell exposure.

**Figure 3:**
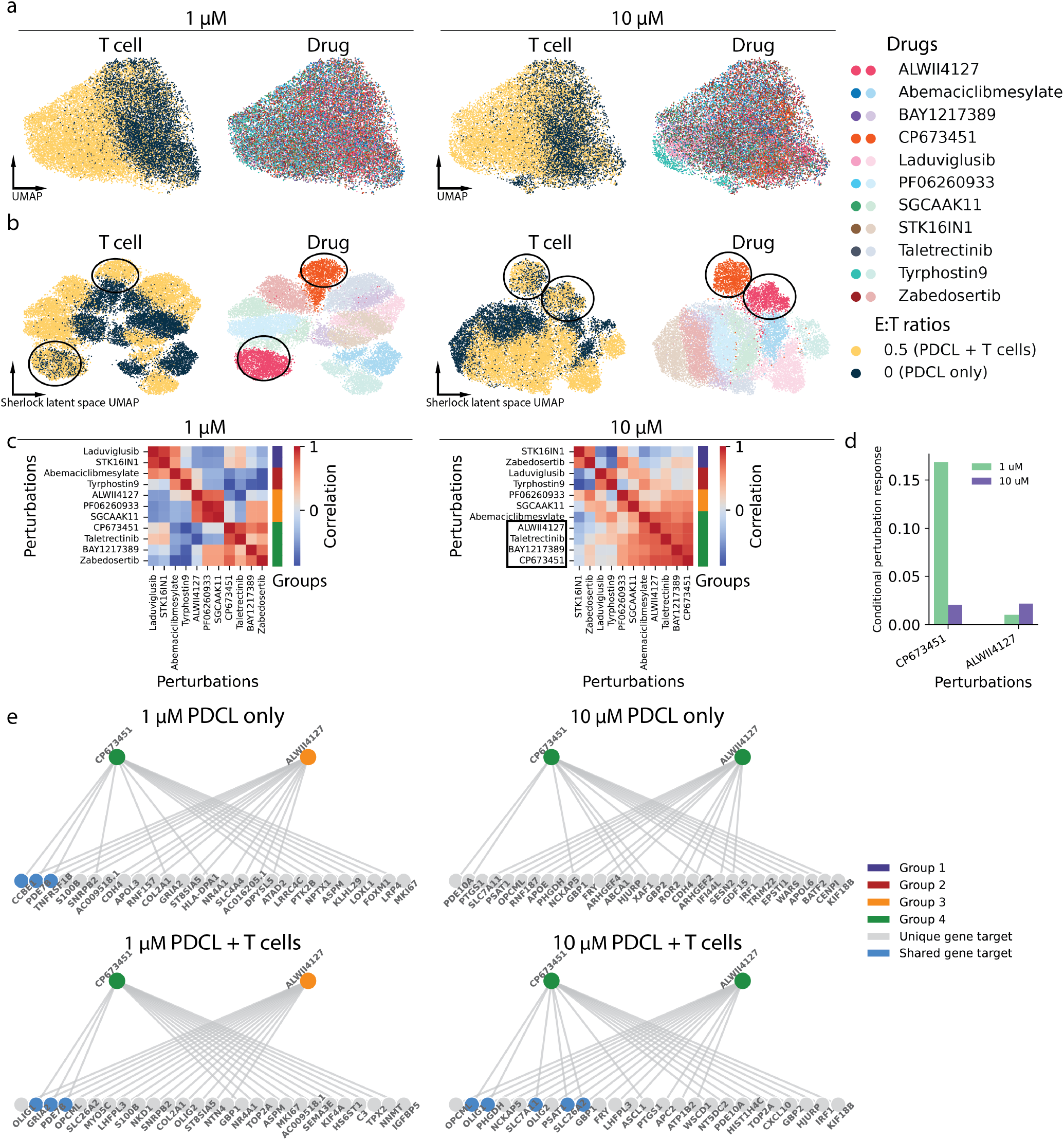
SHERLOCK captures context-dependent drug responses and reveals shared transcriptional programs. (a) UMAP of 11 drug perturbations at 1 *µ*M and 10 *µ*M, colored by T cell co-culture condition (0.5T vs NoT) and drug identity. (b) UMAP of SHERLOCK latent representations for 11 drug perturbations at 1 *µ*M and 10 *µ*M, colored by T cell co-culture condition (0.5T vs NoT) and drug identity. (c) Correlation matrices of perturbation embeddings at each concentration show increased similarity between CP673451 and ALWII4127 at higher concentration, indicating convergence of transcriptional responses. (d) Quantification of condition-specific effects (separation of drug-treated cells by T-cell co-culture condition) using a two-group silhouette score computed in latent space. Positive values indicate strong separation between T cell conditions, whereas values near zero indicate overlap. (e) Causal estimation of downstream genes for drugs CP673451 and ALWII4127 under control (NoT) and T cell co-culture (0.5T) conditions. Increased overlap in downstream target genes is observed under T cell exposure at higher concentrations.

Consistent with Shi et al. [25], CP673451 (CP, a PDGFRA inhibitor) and ALWII4127 (AL, an EPHA2 inhibitor) exhibited concentration-dependent behavior, with their inferred perturbation effects becoming more similar at higher concentrations **(Fig. 3b, c, Fig. S4a)**. Prior work showed that both compounds attenuate the transcriptional response induced by cytotoxic T cell exposure, including blocking the induction of gene programs associated with immune evasion, with ALWII4127 acting across 1–10*µ*M and CP673451 showing the strongest effect at 10*µ*M. This condition-dependent change in drug response is likely the basis by which these compounds enhance T-cell-mediated killing in patient-derived glioblastoma neurospheres [25]. To formally quantify the context-dependence for each drug perturbation, we calculated the separation between cells exposed to different T-cell conditions using a two-group silhouette score in the SHERLOCK latent space **(Fig. 3d)**. Higher scores indicate stronger separation of the drug response by T-cell condition, whereas values near zero indicate greater overlap between conditions. CP and AL showed among the lowest conditional perturbation response scores across the compound panel, indicating convergence of their latent transcriptional responses across T-cell conditions and consistent with drug-mediated modulation of T cell context-dependent gene expression **(Fig. 3b, Fig. S4b)**. Further supporting this convergence, analysis of significant downstream transcriptional effects showed the strongest similarity between CP and AL at 10*µ*M in the presence of T cells **(Fig. S5)**.

Importantly, SHERLOCK further enables causal estimation of downstream transcriptional effects by applying each perturbation as an intervention to an inferred baseline state and decoding the resulting potential outcome. We therefore asked whether the condition-dependent convergence of CP and AL was reflected in their inferred downstream gene programs. SHERLOCK revealed increased overlap between the top 15 genes with the strongest inferred downstream effects of CP and AL under T-cell exposure, particularly at higher concentrations **(Fig. 3e)**. By comparison, a control compound (STK16IN1) showed different concentration-dependent overlap **(Fig. S4c)**.

The shared downstream target genes comprise both glioma lineage/state regulators and immune-response genes. Specifically, *OLIG1* and *OLIG2* are transcription factors associated with oligodendrocyte progenitor-like (OPC-like) glioma cell states [36], while *PHGDH* is associated with metabolic programs in glioblastoma, and *SLC26A2* has been implicated in tumor-cell stress and treatment-response phenotypes [37, 38, 39]. In contrast, *GBP1* is an interferon-inducible effector that has been linked to enhanced antitumor immunity through IFN-*γ* signaling and recruitment of T cells [40, 41]. These two drugs significantly upregulated all shared target genes except **GBP1**, which was significantly downregulated. GBP1 has been described as a “double-edged sword” in cancer, with context-dependent roles as either a tumor suppressor or a pro-survival and pro-tumorigenic factor **(Fig. S6)** [42]. Overall, these results indicate that CP and AL converge under T-cell exposure on downstream transcriptional programs spanning glioma cell-state and interferon-responsive processes, providing a potential transcriptional basis for their previously observed effects on T-cell-mediated tumor killing.

### 2.5 SHERLOCK models combinatorial perturbations as compositional latent interventions

To evaluate the ability of SHERLOCK to model the joint transcriptional effects of perturbations, we applied it to the combinatorial CRISPRa screen of Norman et al. [43], which profiled 105 single and 131 double gene activations in K562 cells (111,255 cells, 3,270 genes after filtering). K562 is a chronic myeloid leukemia cell line derived from blast crisis and can be driven toward erythroid, megakaryocytic, or myeloid fates [44, 45], and the library targets regulators of these fates, MAPK signaling, and the cell cycle. Of the 88 double perturbations with expert genetic-interaction (GI) labels from the original study, 86 retained both genes after filtering and were included for evaluation.

SHERLOCK does not learn an independent embedding for each combination *A*+*B*. Instead, the combined latent shift is constructed from the two constituent single-gene shifts, each scaled by a learned factor between 0 and 2, allowing its contribution to be attenuated or amplified but not reversed, together with a learned non-linear interaction residual (**Methods**). Training is initialized from a purely additive combination, with deviations from additivity penalized so that non-linear interactions are learned only when supported by the data. By reusing the constituent perturbations’ embeddings, gates and covariance structure, SHERLOCK can also predict unseen combinations. These predictions represent compositional extrapolations rather than identified counterfactuals, as the combinations themselves were not observed during training (**Methods**).

In the latent space, double perturbations were organized relative to their constituent single perturbations, with many occupying intermediate positions between them **(Fig. 4a)**. The learned perturbation structure was reproducible across runs with each run’s perturbation correlation matrix correlated with the mean of the other four (*r* = 0.788 ± 0.021), whereas the perturbation effect on latent factors, measured through coactivation of gates due to permutability of latent factors, were less stable (*r* = 0.494 ± 0.056), indicating that reproducibility primarily reflects relationships among perturbations rather than individual latent gates **(Fig. S7)**. Consistent with the recovered perturbation structure, the most strongly coupled perturbations were paralogous genes or genes with related biological functions, specifically, *ETS2* /*MAPK1* (*r* = 0.89 ± 0.02), the CIP/KIP inhibitors *CDKN1A*/*B*/*C* (*r* = 0.79–0.89 across the three pairs, SD ≤ 0.12) [46], *CEBPA*/*CEBPB* (*r* = 0.87 ± 0.06) [47], the two p38 MAP2Ks *MAP2K3* /*MAP2K6* (*r* = 0.82 ± 0.03) [48], and *IGDCC3* /*PRTG* (*r* = 0.81 ± 0.04).

**Figure 4:**
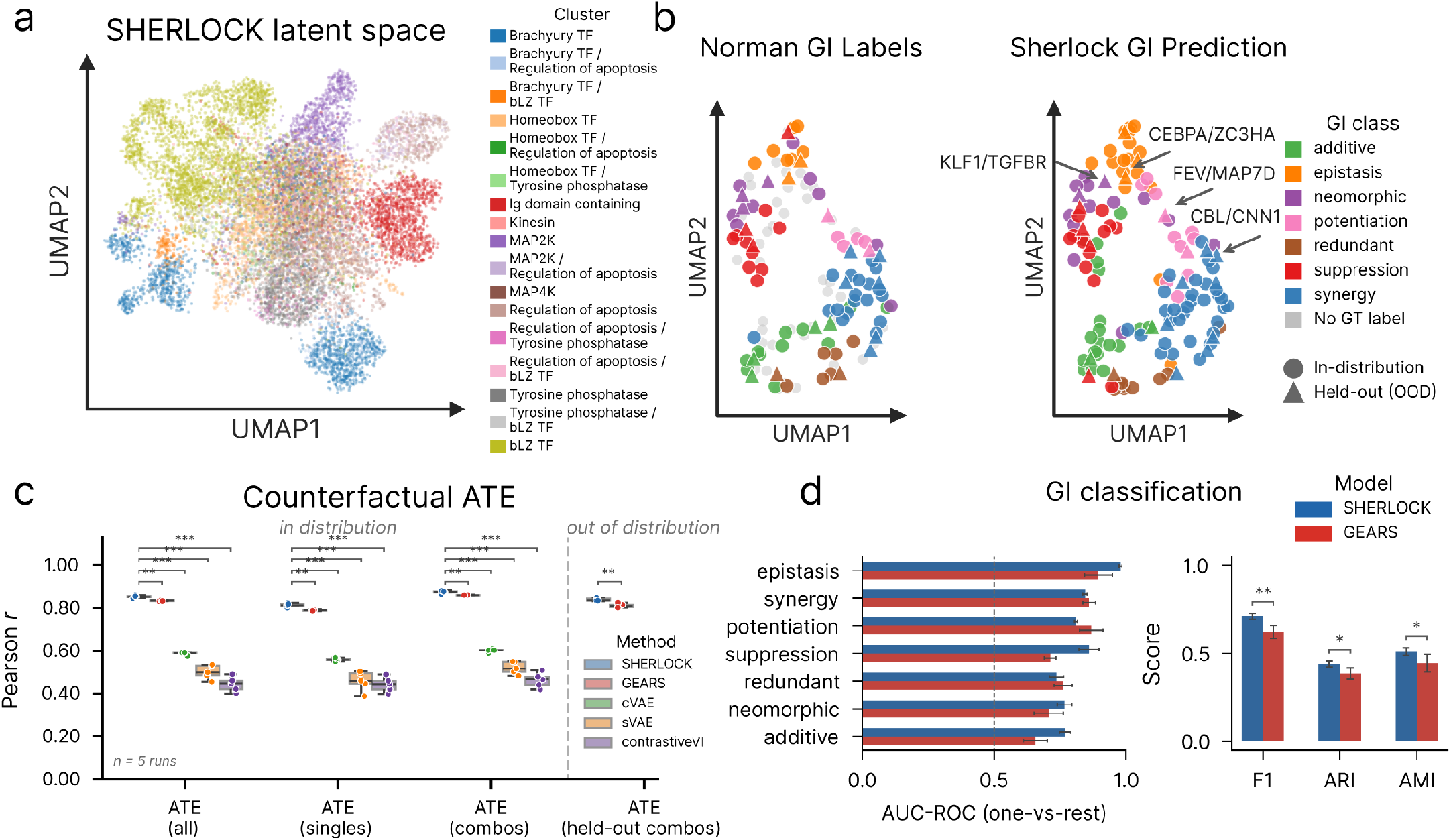
SHERLOCK models combinatorial perturbations and recovers genetic-interaction categories in the Norman CRISPRa screen [43]. All statistics are over *n* = 5 seeds per method (Welch’s *t*-test, Benjamini–Hochberg corrected). (a) SHERLOCK latent space, colored by the perturbation modules annotated in (Supp. Table 6 of) the original study. (b) Embedding of the 11 genetic-interaction metrics, colored by Norman labels (left) and SHERLOCK prediction in observed mode (right); circles, seen pairs; triangles, held-out pairs. (c) Counterfactual average treatment effect (Pearson *r*, predicted versus observed effects) across all perturbations, singles, seen and held-out combinations. Held-out ATE is undefined for methods that treat a combination as an atomic label (cVAE, sVAE, contrastiveVI, scGen). (d) Genetic-interaction classification over the 86 labeled pairs (observed mode, synergy subtypes merged): per-class AUC-ROC (left) and weighted F1, ARI and AMI (right).

To test SHERLOCK’s ability to predict unseen perturbation combinations, we held out 22 of the 86 labeled pairs, stratified by GI class, and compared their predicted and observed average treatment effects (ATEs) **(Fig. 4c)**. Across five seeds, SHERLOCK achieved a Pearson correlation of *r* = 0.852 ± 0.004 across all combinations and *r* = 0.836 ± 0.005 on held-out combinations, compared with *r* = 0.833 ± 0.001 and *r* = 0.810 ± 0.008, respectively, for GEARS [11] (BH-adjusted *q <* 0.003). In contrast, generative baselines that model each combination as a distinct perturbation showed lower overall performance (cVAE, *r* = 0.498; contrastiveVI, *r* = 0.496; sVAE, *r* = 0.428) and could not generalize to unseen combinations.

### 2.6 SHERLOCK recovers genetic-interaction structure from combinatorial perturbation effects

We next evaluated whether the modeled perturbation effects recapitulated the interaction categories defined by Norman et al.[43], which were not used during model training. As performed in Norman et al., we linearly regressed the effect of each double-perturbation on the effects of its two constituent single-perturbations and derived 11 metrics quantifying magnitude, dominance, fit and similarity, which were used to assign each pair to one of seven interaction classes across 5 independent SHERLOCK runs. The same procedure was applied to effect vectors obtained from SHERLOCK and GEARS and, as a reference, to measured expression.

Across the 86 labeled pairs, SHERLOCK outperformed GEARS on weighted F1 (0.712 ± 0.017 vs 0.623 ± 0.035), ARI (0.440 ± 0.016 vs 0.387 ± 0.033) and AMI (0.512 ± 0.022 vs 0.446 ± 0.050; BH-adjusted *q* = 0.008–0.040; **Fig. 4d**), with the largest class-specific gains for approximately additive (AUROC 0.770 ± 0.021 vs 0.657 ± 0.045) and suppression (0.860 ± 0.038 vs 0.713 ± 0.022) interactions. Measured expression performed similarly to SHERLOCK (weighted F1 0.700 ± 0.012 across five regression seeds; Welch’s *t*-test, *p* = 0.24), with higher performance on neomorphic pairs and lower performance on approximately additive pairs (**Fig. S9**). Because Norman’s interaction categories were defined from measured expression, this analysis tests how faithfully each model preserves the interaction structure present in the observed data. On the 22 held-out pairs, whose effects were predicted compositionally from their constituent single perturbations, SHERLOCK and GEARS showed comparable performance (weighted F1 0.692 0.047 vs 0.696 0.120, *q* = 0.95; **Fig. S8**). Representative pairs from each interaction class are shown in **Fig. 5a** and **Fig. S10**.

**Figure 5:**
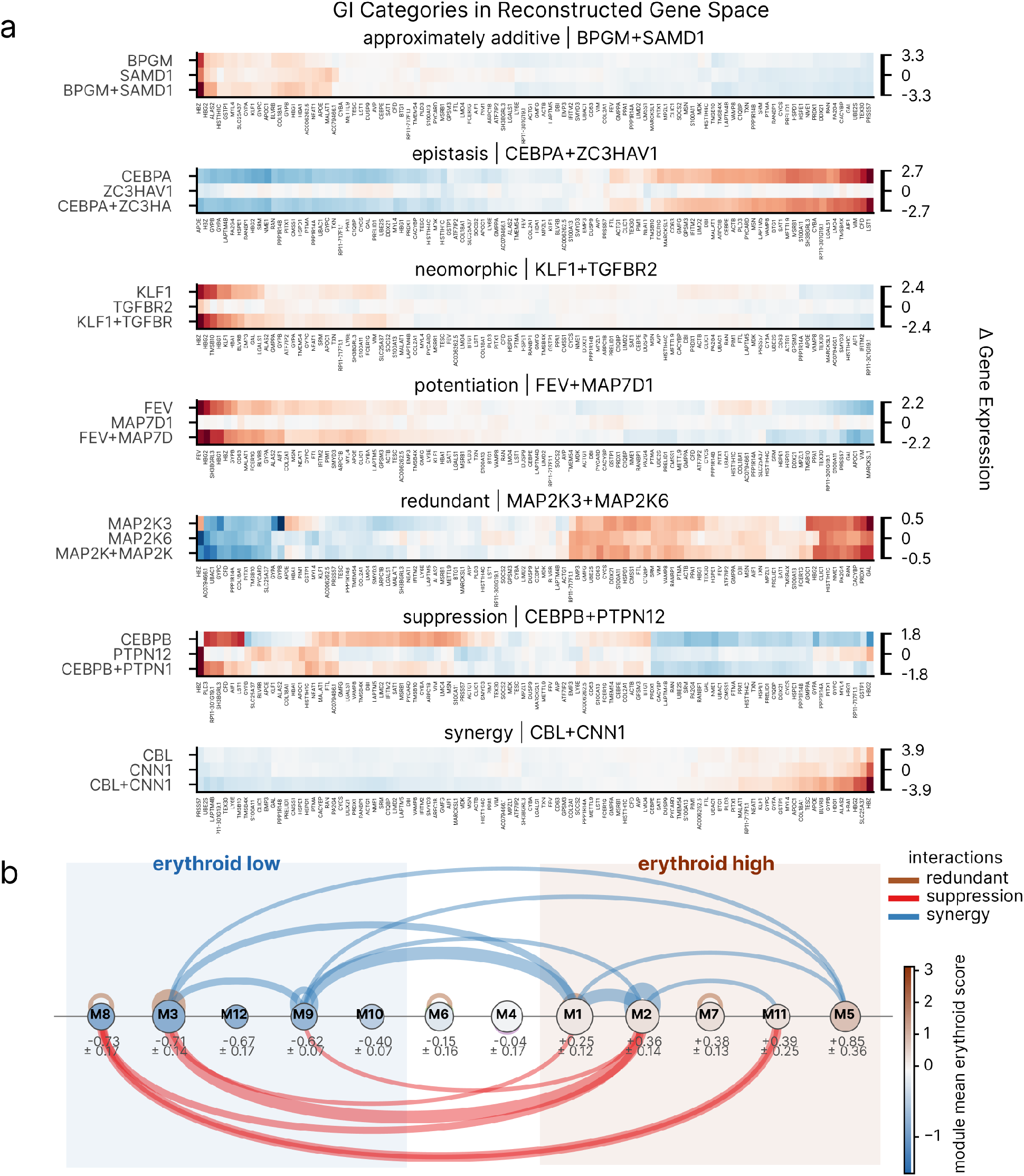
Genetic interaction classes reveal distinct transcriptional effects and lineage-associated module relationships. (a) Reconstructed responses to the two constituent single perturbations and their combination for a representative pair per interaction category (100 highest-variance genes). (b) Each labeled pair is drawn as an arc between the consensus modules of its two parents. Modules run from erythroid low on the left to erythroid high on the right, with their mean observed erythroid score and its standard error over member perturbations printed below each one; node color repeats that score (color bar) and node size scales with the number of members. Arc color indicates the interaction class including redundant, suppression, and synergy.

A regression on expression profiles describes how a combination relates to its constituent perturbations, but not how that relationship is represented by the model. Although the model is never trained on interaction labels, its structured parameters indirectly contain information about how two perturbations combine, thus allowing interaction classes to also be inferred from the fitted model parameters, rather than from the expression profile alone. We therefore summarized every labeled pair with seven descriptors of perturbation similarity, latent-factor sharing, single-perturbation scaling and the non-additive residual (**Methods**, Section 4.8.3). A decision list over these quantities, aggregated across 5 runs to obtain a singular parametrization and applied in a fixed order, classified the labeled pairs with an overall accuracy of 0.77 (and a balanced accuracy of 0.78; with recall ranging from 0.50 for neomorphic to 0.92 for suppression and 1.00 for the six potentiation pairs) (**Fig. S11a,b**), showing that the fitted model parameters contain substantial information about the annotated interaction classes. Because the rule thresholds were selected using the same 86 labeled pairs, this analysis measures how well SHERLOCK’s learned parameters separate known interaction classes (Additive, Redundant, Synergy, Potentiation, Epistasis, Suppression, and Neomorphic), rather than the ability of these rules to classify unseen combinations.

The rule for neomorphic interactions rests on two properties of the fitted model (**Fig. S11c,d**). First, once decoded, 59-69% of the effect added by the interaction term across the annotated classes lies outside the span of the two constituent single-gene perturbations (also referred to as parents). Second, across all composable pairs, a median of 33% of this decoded effect arose from latent factors that neither single perturbation uses, and this component was only weakly correlated with the parents’ single perturbation gates (|Spearman| ≤ 0.41). The interaction term can therefore move the cell in a direction neither single perturbation reaches, which is what defines a neomorphic combination.

### 2.7 SHERLOCK links genetic interaction structure to erythroid-associated transcriptional programs

To connect the learned interactions to biological function, we constructed 12 consensus perturbation modules by co-clustering the 105 single perturbations across the five runs and compared these modules with the functional clusters of Norman et al. [43]. Nine of the 20 reference clusters were more enriched in one module than expected for random modules of the same sizes (79% vs 41% of members; *q* = 0.004-0.032; **Fig. S12a**). In addition, the modules recovered biologically coherent groupings of perturbations, including a module of tyrosine phosphatases (6/8 in M2), members of the bZIP transcription factors and granulocyte group (all in M3) and CDK inhibitors (all in M7).

To characterize the transcriptional responses associated with these modules, we calculated canonical erythroid and granulocytic marker scores from measured expression changes relative to unperturbed cells (**Methods**). Erythroid and granulocytic scores were negatively correlated (*r* = ™0.31 ± 0.09, *q* = 0.009), and observed erythroid scores were reproducible (*r* = 0.80 ± 0.03). We used the observed erythroid score to organize modules along a continuous transcriptional axis. Placing each module at the mean observed erythroid score of its members, the modules fall into three poles, split at a module-mean erythroid score of ±0.2 with within-module SD ranging from 0.13-1.15 (**Fig. 5b, Fig. S13**).

The erythroid-high pole contains M2, which included four members of the erythroid phenotype group described by Norman et al. (*CNN1, PTPN9, PTPN12, UBASH3B*), M5 (erythroid +0.85 ± 0.36, megakaryocytic +0.50; *FEV, IKZF3*), M7 (G1-arrest group *CDKN1A*/*B*/*C*), and M1 (+0.25 ± 0.12) and M11 (+0.39 ± 0.25), with the latter two lying within one standard error of the threshold defining the poles. M4 and M6 were intermediate. At the erythroid-low pole, M3 had a high granulocytic marker score (+1.12) and contained C/EBPs and *SPI1*, regulators of granulocytic and monocytic differentiation [47, 49]. By comparison, M8, M9, M10 and M12 (erythroid −0.40 to −0.73, SEM ≤ 0.17; M8 containing *MAPK1* /*ETS2*) had low erythroid scores without a comparably high granulocytic or megakaryocytic signature (granulocytic ≤ +0.36).

To examine how these programs were represented in latent space, we aligned factors across runs (using Hungarian matching; *q* = 0.0002 consistency per run; **Fig. S14**). One reproducible latent factor (F13; cross-run signature similarity | cos | = 0.86 ± 0.08, used by 87 ± 5 of the 105 perturbations) had the highest erythroid program score (+5.1 ± 0.8, compared to +2.8 ± 0.6 for the next factor), with *IKZF3, DUSP9* and *TP73* showing strongest positive association, and the C/EBPs as its strongest negative association. A second latent factor (F12) captured the alternative lineage-associated programs (granulocytic score +1.5 ± 0.3; erythroid 1.3 ± 2.2), indicating greater cross-run consistency in the myeloid component (cross-run similarity 0.79 ± 0.10): with C/EBPs and *MAPK1* /*ETS2* exhibiting strongest positive association on the granulocytic end, and *CNN1* and *PRTG* on the erythroid end (**Fig. S15**). This organization is consistent with the established antagonism between erythroid and myeloid regulators [50, 47] and MEK/ERK signaling altering K562 differentiation [51].

We next examined whether these module-level transcriptional patterns were associated with genetic-interaction classes (**Fig. 5b, Fig. S13**). Within the annotated panel, all 8 redundant pairs combined perturbations within the same module (*q* = 4 × 10^*−*5^), with 5 in the erythroid-low pole, including the C/EBP paralogues (*q* = 0.003). In contrast, none of the 12 suppressive pairs lay within a module (*q* = 0.035) and all 12 combine an erythroid-high with an erythroid-low module (*q* = 2 × 10^*−*5^), setting the C/EBPs and *MAPK1* /*ETS2* against the erythroid phosphatases, *PRTG*/*IGDCC3* and *LYL1*. Independent of module assignments, 11 of the 12 pairs also had constituent single perturbations with opposite-signed erythroid scores (*q* = 0.003). Synergy was enriched within the erythroid-high pole (11/24; *q* = 0.035) and additivity rarely spanned the poles (2/15; *q* = 0.040). However, these latter associations were sensitive to the module classification threshold (*q* = 0.18 and *q* = 0.32).

Module membership did not uniquely determine interaction class. Nine pairs spanning erythroid-high and erythroid-low modules were annotated as synergistic, including 7 dissimilar-phenotype synergies. These pairs mostly involved forkhead and homeobox transcription factors whose constituent single perturbations had similar erythroid scores despite belonging to modules in different groups. Six of the nine pairs had positively correlated perturbation representations (*ρ >* 0.2, compared to 0/12 suppression pairs) and decoded joint-effect magnitudes exceeding the additive prediction. In contrast, pairs more widely separated along the erythroid-score axis had perturbation correlations closer to zero. These examples illustrate how module-level differences can coexist with shared structure in the learned perturbation representations. Overall, these analyses connect SHERLOCK’s perturbation modules and latent factors to lineage-associated transcriptional programs and reveal associations between this organization and genetic-interaction classes.

## 3 Discussion

In this work, we introduce SHERLOCK, a structured generative framework that learns interpretable perturbation representations and estimates the causal effects of perturbations on downstream gene expression in single-cell perturbation data. By jointly modeling baseline cellular states, partially shared perturbation mechanisms, and their dependence on biological condition, SHERLOCK organizes genetic and pharmacological perturbations according to shared response programs and supports comparisons across biological conditions. Across Perturb-seq, imaging-based spatial perturbation data, and sci-Plex datasets, SHERLOCK recovers biologically coherent perturbation groups spanning CRISPRi, CRISPRa, CRISPR knockout, and drug treatments. These findings support the applicability of structured perturbation modeling across diverse measurement technologies and intervention modalities.

A central contribution of SHERLOCK is its integration of interpretable representation learning with an explicit structural causal formulation. Sparse perturbation gates, constraints informed by basal-to-perturbed gene expression differences, and a low-rank perturbation covariance structure restrict the latent representation while capturing relationships among interventions. Our theoretical analysis specifies the SCM view of the model and its induced causal estimand. We establish identifiability within a restricted model class under explicit assumptions of causal validity and treatment coverage, identifiability of the noisy observation model, common affine alignment of latent coordinates, and specified shared-noise coupling across interventions. The theory extends to combinatorial perturbations represented in the observed data under the corresponding assumptions. Across independent training runs, estimated counterfactual transcriptional effects and relationships among perturbations were more reproducible than gate co-activation patterns. This supports the practical stability of these outputs, even though theoretical assumptions do not necessarily hold in practice.

Our findings further emphasize that perturbation effects depend on cellular context. In the T cell co-culture setting, CP and AL showed increasingly similar inferred effects at higher concentrations, alongside reduced separation between transcriptional states with and without T-cell exposure. These observations complement prior experimental evidence that both compounds enhance T cell–mediated tumor killing [25]. By comparing perturbation responses with condition-matched control populations, SHERLOCK helps distinguish perturbation-induced transcriptional responses from those induced by changes in the cellular context. The resulting shared gene programs suggest that these compounds may act through both modulation of tumor-cell immune-response programs and tumor-state remodeling. Previous studies have associated tumor-state transitions with antitumor drug sensitivity [52] and T-cell dysfunction or exhaustion in glioblastoma [53]; our results motivate further investigation of how these programs influence susceptibility to T cell–mediated killing. In a complementary kinase-focused CRISPRi/a screen, SHERLOCK also identified established regulators, including JAK1 and AXL, and nominated MAP4K3 and PIM2 as candidate regulators of tumor immune adaptation.

The combinatorial analyses extend this framework from characterizing individual perturbations to predicting joint responses. By composing effects of constituent single perturbations, SHERLOCK predicted held-out combinations and improved average-treatment-effect prediction relative to previous methods. Associations between perturbation modules, interaction classes, and erythroid-associated expression further connect the learned perturbation structure to biologically relevant variation in joint responses. These findings provide hypotheses about how combinations reshape transcriptional programs, while functional differentiation and specific regulatory mechanisms require independent validation.

Several limitations provide directions for future work. Counterfactual interpretation depends on the validity of the assumed causal structure and the extent to which the learned representation satisfies the identifiability conditions. Moreover, transcriptional programs and candidate regulators inferred computationally require experimental validation to establish their mechanistic roles. Incorporating chromatin accessibility or protein measurements could strengthen the connection between latent mechanisms and molecular regulation, while explicit temporal and spatial modeling could capture dynamic responses and interactions between cells. These extensions could advance SHERLOCK as a framework for investigating context-dependent perturbation mechanisms, prioritizing therapeutic targets, and investigating combination strategies in immuno-oncology.

## 4 Methods

### 4.1 Notations and Minibatch Construction

Let *G* denote the number of genes, *d* the latent dimension, and *P* the number of modeled perturbation labels, excluding the separately supplied non-targeting control (NTC) observations. Each perturbed cell has one label *p*_*n*_ ∈ {1, …, *P*}. A minibatch contains *B* perturbed count vectors 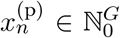 and a row-aligned array of *B* control count vectors 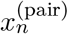 . For control reconstruction, we remove duplicate rows from the paired control array, leaving *M* ≤ *B* distinct control count vectors, denoted by 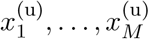. We then form a combined batch of size *T* = *M* + *B* by placing these *M* control observations before the *B* perturbed observations. Thus, row *i* corresponds to control observation *i* for *i* = 1, …, *M*, and row *M* + *n* corresponds to perturbed observation *n* for *n* = 1, …, *B*. Each observation in this combined batch has two latent variables, 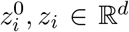. Any condition dependence of the background posterior comes from the supplied control expression profiles.

### 4.2 Generative model

We structure perturbation effects through a low-rank row covariance and learned coordinate-wise modulation. This parameterization encourages shared structure but does not by itself guarantee identifiability of the latent representation.

#### 4.2.1 Low-rank perturbation covariance

Let *P* denote the number of perturbations, *d* the latent dimension, and *r* ≤ *P* the rank of the low-rank covariance component. Write *I*_*k*_ for the *k* × *k* identity matrix. Let *U*_cov_ ∈ ℝ^*P*×*r*^ be a learned unconstrained matrix and *l* ∈ ℝ^*r*^ a learned vector of log-scales. Denote the reduced QR factors of *U*_cov_ by *Q* ∈ ℝ^*P*×*r*^ and *R* ∈ ℝ^*r*×*r*^, so that

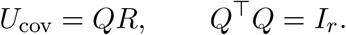

We define the positive scale vector *s* ∈ 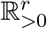 by *s*_*k*_ = exp(*l*_*k*_). On each forward call, we adjust the signs of the columns of *Q* using the diagonal entries of *R*, then jointly reorder the columns of *Q* and entries of *s* by decreasing scale. The low-rank covariance factor *L* ∈ ℝ^*P*×*r*^ is then

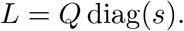

Let *σ >* 0 be a learned isotropic standard deviation. Define the perturbation covariance matrix Σ_*P*_ ∈ ℝ^*P*×*P*^ and its lower-triangular Cholesky factor *C*_*P*_ ∈ ℝ^*P*×*P*^ by

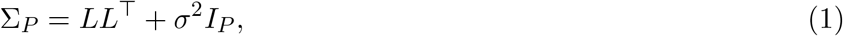

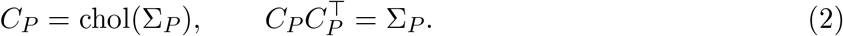

The term *σ*^2^*I*_*P*_ ensures that Σ_*P*_ is positive definite. Let *ρ* ∈ R^*P*×*d*^ be a latent matrix with rows *ρ*_*p*_ ∈ ℝ^*d*^, and let *b* ∈ ℝ^*P*×*d*^ collect their prior means *b*_*p*_ R^*d*^. We assign independent Gaussian priors to these rows:

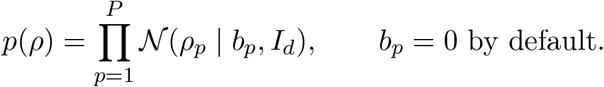

The perturbation-effect matrix *A* ∈ ℝ^*P*×*d*^ is defined by

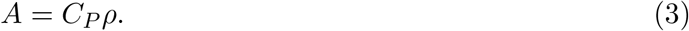

For *j* = 1, …, *d*, let *A*_:*j*_ and *b*_:*j*_ denote the *j*th columns of *A* and *b*. Conditional on *L, σ*, and *b*, the columns of *A* are independent, with

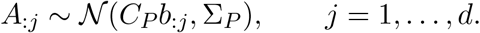

Thus *A* has a matrix-normal prior with mean *C*_*P*_ *b*, row covariance Σ_*P*_, and column covariance *I*_*d*_; its mean is zero when *b* = 0. The prior on *A* is induced by the independent Gaussian prior on *ρ* and the deterministic transformation in Eq. (3).

#### 4.2.2 Latent Coordinate Modulation and Gradient

For each perturbation and latent coordinate, let *α*_*pj*_ be a learned logit and draw independent *u*_*pj*_ ∼ Uniform(0, 1). Write sigm(*a*) = (1 + *e*^*−a*^)^*−*1^, we have

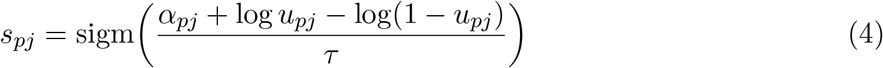

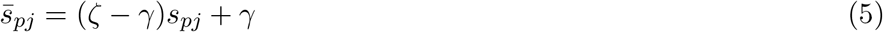

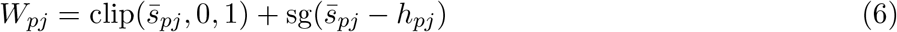

where sg denotes stop-gradient, *γ* = ™0.1, *ζ* = 1.1, and *τ >* 0. Away from clipping boundaries, the backward gradient rule is

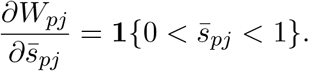

Consequently, this is a stretched Concrete variable with a clipped surrogate gradient, rather than a standard hard-clipped HardConcrete forward variable. It can reverse or amplify a coordinate’s shift. A single *P × d* sampling is shared across observations in each model evaluation. The deterministic gate used during inference replaces *s*_*pj*_ with sigm(*α*_*pj*_*/τ*) and has the same unclipped forward form. We also compute

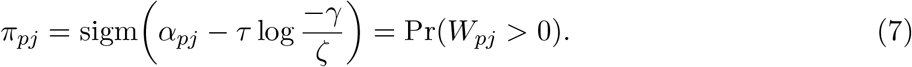

as the usage surrogate that shows up in the regularizers formulation.

#### 4.2.3 Background state and latent transition

For a single perturbation, define *δ*_*p*_ = *W*_*p*_ ⊙ *A*_*p*_. Each unique control row has shift Δ_*i*_ = 0, while a perturbed row has 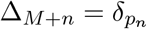 . We model the base cell

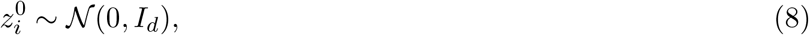

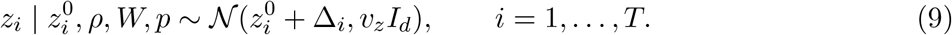

The scalar *v*_*z*_ *>* 0 is the variance scalar and has default value 1. For control rows, the conditional mean of *z*_*i*_ is 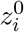.

#### 4.2.4 Monte Carlo background sampling

Because 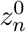 for a perturbed cell is the encoding of a randomly drawn NTC cell rather than of 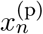 itself, each minibatch supplies a fresh background draw for every perturbed cell, so that over training the expectation in the objective is taken over the control population as a whole rather than over any one background state. Sampling with replacement means one NTC cell may be paired with several perturbed cells within a batch. The duplicated rows are exact copies, so before evaluating the background reconstruction term we reduce the batch’s NTC rows to the distinct cells actually drawn; this keeps the NTC likelihood from being counted once per pairing and keeps the cost of the background decode independent of the perturbed batch size. The same reduction is applied in the model and in the guide so that the two agree on the number and order of background rows.

#### 4.2.5 Negative binomial decoder and auxiliary classifier

Let 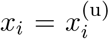 for control rows and 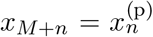 for perturbed rows, and let 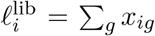 be the observed library size. The shared decoder computes

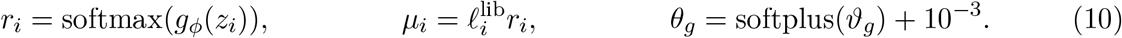

Define the NB logits *η*_*ig*_ = log(*µ*_*ig*_ + *ϵ*) − log *θ*_*g*_. We have

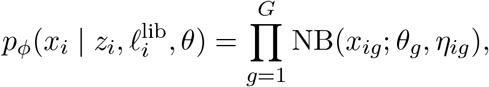

where *ϵ* is a small constant and the second and third arguments specify total count and logits. Both control and perturbed counts are decoded from *z*_*i*_, not from 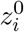. However, for control rows, the conditional mean of *z*_*i*_ is 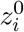.

The classifier *h*_*ψ*_(*z*_*M*+*n*_) ∈ ℝ^*P*^ is applied only to the perturbed rows. Its categorical cross-entropy loss is

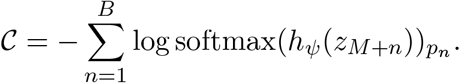

The classifier is an auxiliary supervised loss, since the perturbation labels already determine the transition shifts.

### 4.3 Amortized Variational inference

Let *E*_0_ denote the background encoder and its mean/log-variance head, and *E*_*z*_ the separate perturbed-cell encoder and head. For unique controls,

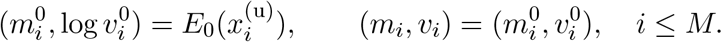

For perturbed rows,

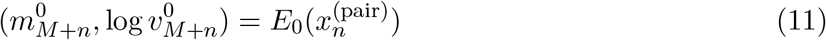

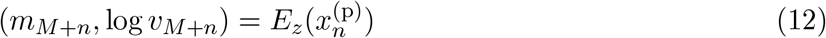

The variational family factorizes as

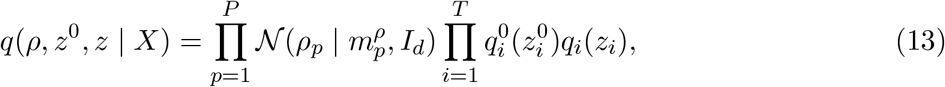

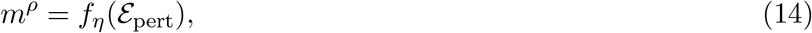

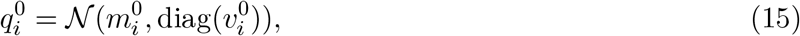

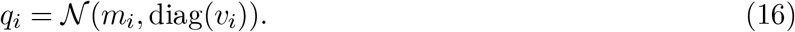

Here _pert_ ∈ ℝ^*P*×*d*^ is a learned embedding table and *f*_*η*_ is a learned network. The covariance of *q*(*ρ*) is fixed to identity. For controls, *q*_*i*_ and *q*^0^ share their parameters but are sampled at distinct sites, so they are independent given the encoder outputs, not the same random variable. Repeated paired backgrounds also lead to separate latent samples for their corresponding perturbed rows. Both encoders receive gradients through their associated likelihood and variational terms; the perturbed encoder also receives classifier gradients. Each expression encoder has two width-128 linear layers, each followed by LeakyReLU and LayerNorm, and a linear head returning 2*d* values. The decoder has one width-128 hidden layer with LeakyReLU. The classifier has two width-128 hidden layers with LeakyReLU. The perturbation-mean network has two *d*-dimensional linear layers separated by LeakyReLU.

### 4.4 Variational objective and regularization

We describe the summed Minibatch objective under standard ELBO inference. Because likelihood weights, auxiliary classification, and additional factors are present, we call the complete expression a regularized variational objective. Let *U* collect the independent uniforms used for *W*, and suppress the fixed library-size offsets in the likelihood notation. The objective to maximize is

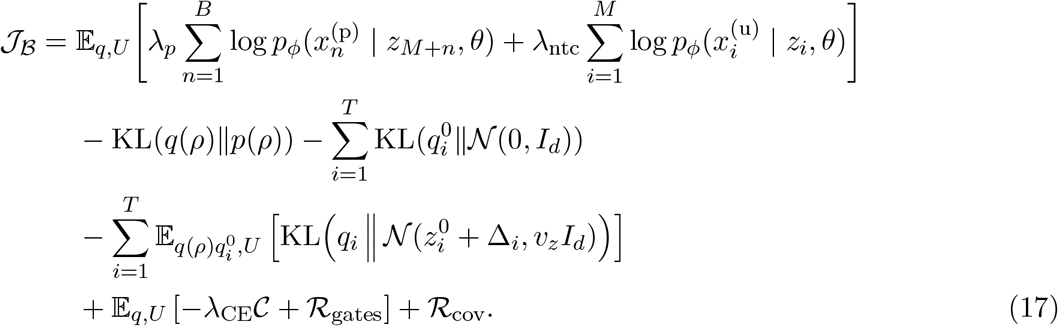

where *C* is the cross entropy of the classifier, *ℛ*_gates_ is the gate regularizer, and *ℛ*_cov_ is the covariance regularizer, for which we give precise definitions later on. The global *ρ* KL and covariance factors occur once per model call; local KL terms include both control and perturbed rows. Priors already represented inside the KL terms are not added again. In particular, there is no additional log *p*(*A L, σ*) or implicit log *p*(*θ*) term. The gate parameters are learned as free parameters. Equation (17) describes the expected forward score; gate regularizer follow the surrogate rule in Eq. (6).

#### Covariance Regularizer

We add a log-factor penalty

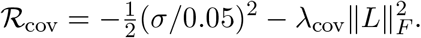

It is applied to the point-estimated positive parameter *σ* and factor *L*; *σ* is initialized to 0.05.

#### Gating regularizers

With *ϵ* = 10^*−*8^, define

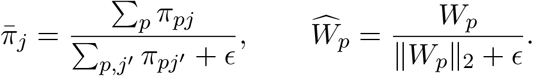

The gating regularizer is

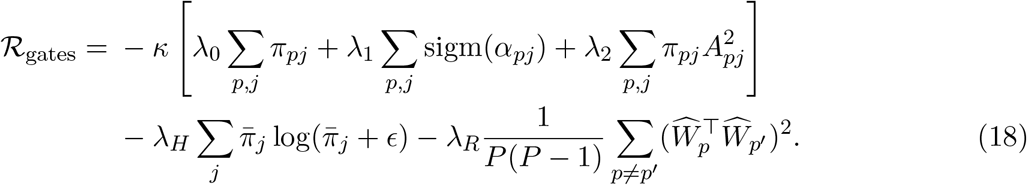

The row-repulsion term is omitted when *P* = 1. *κ* is a scaling factor hyperparameter. The entropy and row-repulsion terms remain active whenever their coefficients are nonzero. In the square bracket, the second term is an *L*_1_ penalty on sigmoid-transformed logits. The third is a usage-weighted quadratic penalty on *A*.

### 4.5 Quantifying condition dependence with a silhouette score

To ask whether a perturbation’s effect depends on the condition it is applied in, we score each perturbation separately in the learned latent space. Taking only the cells carrying that perturbation and labelling them by condition (for example with and without T-cell co-culture), we compute for each cell *i* its mean Euclidean distance *a*(*i*) to the other cells of its own condition and its mean distance *b*(*i*) to the cells of the other condition, and combine them into the silhouette *s*(*i*) = (*b*(*i*) ™ *a*(*i*))*/* max {*a*(*i*), *b*(*i*)} . The perturbation’s score is the mean of *s*(*i*) over all of its cells in the two conditions, and is left undefined when either condition contributes fewer than two cells.

The score lies in [™1, 1]. A high value means the perturbation’s cells separate by condition in latent space, so the perturbation’s effect is reshaped by the condition; a value near zero means the two conditions overlap, so the perturbation acts in the same way in both; negative values indicate the conditions are interleaved. Because the score depends only on relative distances, it is invariant to the scale and orientation of the latent space and is therefore comparable across perturbations and across independently trained runs. Scoring each perturbation on its own cells makes the comparison internal to that perturbation, so differences in how many cells a perturbation has, or in how strong its overall effect is, do not by themselves change the score.

### 4.6 Causal Inference of Perturbation Effect

We describe how the additive SHERLOCK model can be interpreted as a structural causal model (SCM), and distinguish counterfactual quantities defined by that SCM from quantities identifiable from observed data. The motivating query is at level *ℒ*_3_ of the Pearl Causal Hierarchy [15, 14]: *Given that a cell was observed under control with counts x*^(ntc)^, *what would its expression have been under perturbation p?* We first define the SCM and its counterfactual estimand, then state sufficient assumptions for identifiability, and finally describe the approximation implemented by the prediction helpers.

#### 4.6.1 SCMs, Interventions, and Counterfactuals

To begin our discussion, we provide some theoretical background on Pearl Causal Hierarchy and the SCM framework. Under Pearl’s causal framework, structural causal models induce associational, interventional, and counterfactual distributions. We first formally define structural causal models as the following:

##### Definition 1

(Structural Causal Models). *A Structural Causal Model (SCM) is defined as a 4-tuple ℳ* = ⟨*U, V*, F, *P* (*U*)⟩:

1. *Exogenous Variables(U): A set of unobserved background (noise) variables determined by factors outside the model*
2. *Endogenous Variables(V): A set of observed or latent variables determined by other variables in the model, more explicitly by the set U* ∪ *V*
3. *Causal Mechanisms (ℱ): A set of functions* {*f*_1_, *f*_2_, …, *f*_*n*_} *such that each f*_*i*_ *maps from the respective domains of U*_*i*_ ∪ *Pa*(*i*) *to V*_*i*_, *where U*_*i*_ ⊆ *U and Pa*(*i*) ⊆ *V* \ {*V*_*i*_}, *and is given by*

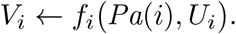
4. *Joint Noise Density(P* (*U*)*): A joint probability distribution over the exogenous variables*.

With this definition, we can analyze the process of SCM inducing various layers of Pearl Causality. Trivially, the mechanism set *ℱ* and the prior *P* (*U*) together induce the observational distribution *P* (*V*) over the endogenous variables. But in order to explain how interventional and counterfactual distributions are induced, we need to define the process of performing an intervention:

##### Definition 2

(Interventions and Interventional SCM). *An intervention, denoted by the do*(·)*-operator, modifies the SCM into a sub-model ℳ*_*x*_ = ⟨*U, V*, F_*x*_, *P* (*U*)⟩, *where for X* ⊆ *V*

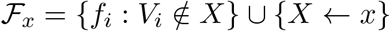

*Such a sub-model is called an interventional SCM*.

From this definition, we can write out the *interventional distribution* induced by interventions on an SCM for each *Y* ⊆ *V* as:

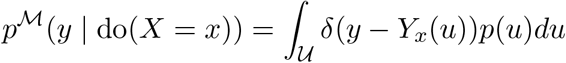

where *Y*_*x*_(*u*) is the *potential outcome* of *Y* under intervention do(*X* = *x*).

Now, if we consider a set of potential outcomes I = {*Y*_*x*_, …, *Z*_*w*_} governed by a set of interventional submodels M = {ℳ_*x*_, …, *ℳ*_*w*_} all induced from the base SCM ℳ, the *joint counterfactual distribution* over I is defined by marginalizing out the underlying exogenous background variables *U* shared across all submodels, i.e.

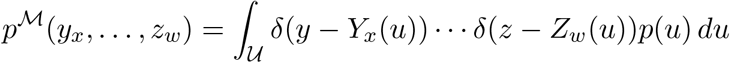

Crucially, this joint distribution requires only a *single* integration over the exogenous noise *P* (*U*). This distinguishes it from a product of marginal interventional distributions, where the background noise *u* would be independently drawn for each do(·) operator.

#### 4.6.2 Causal Inference with SHERLOCK

For discrete count outcomes, we write Dirac-delta as indicators. We first consider one fixed experimental context and single perturbations. In this section, *P* denotes the assigned perturbation label, taking values in a finite set *P* that includes the control label ∅. Let *d* be the latent dimension and *G* the number of genes. We condition on fixed fitted weights, including the perturbation-effect matrix *A*, gate matrix *W*, decoder parameters *ϕ*, and gene-wise negative binomial parameters 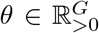 .

Define

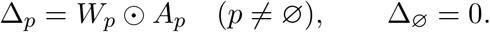

At prediction time, *A* is computed using the posterior mean of the global embedding and *W* is the deterministic gate output. Fix an externally specified decoder library-size offset *l >* 0. All identifiability statements below are on the corresponding fixed-offset population model.

Let *g*_*ϕ*_ : ℝ^*d*^ *→*ℝ^*G*^ be the decoder’s logit map and *ε*_NB_ = 10^*−*6^ its numerical stabilizer. Define the NB mean vector and observation kernel by

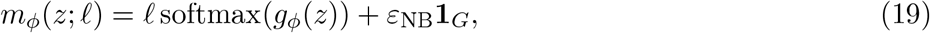

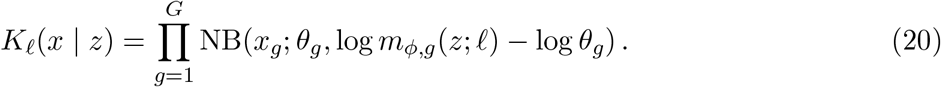

For each gene *g*, let *F*_*g*_(; *z, l*) be the corresponding count CDF and 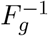 its generalized inverse. To specify a complete SCM, choose the observation mechanism

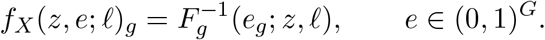

This inverse-CDF choice supplies a particular coupling of observation noise across interventions. The NB likelihood alone does not specify this coupling.

##### Definition 3

(SCM view of SHERLOCK). *Let v*_*z*_ *>* 0 *be the fixed transition variance. Consider mutually independent exogenous variables ϵ*^*P*^ ∼ Uniform(0, 1), *ϵ*^0^ ∼ *N* (0, *I*_*d*_), *ϵ*^*z*^ ∼ *N* (0, *I*_*d*_), *and ϵ*^*x*^ ∼ Uniform((0, 1)^*G*^), *where ϵ*^*P*^ *generates the assignment P through a function f*_*P*_ . *The endogenous variables are V* = {*P, Z*^0^, *Z, X*}, *with structural assignments*

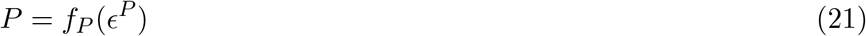

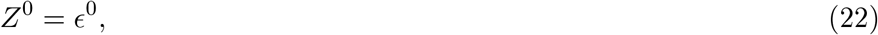

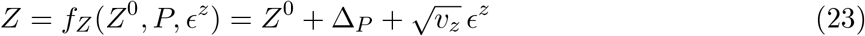

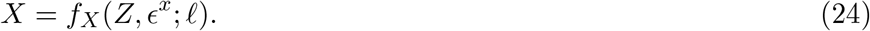

*This defines ℳ*_SH_ = *U, V*,, *P* (*U*), *with U* = (*ϵ*^*P*^, *ϵ*^0^, *ϵ*^*z*^, *ϵ*^*x*^) *and the product noise distribution P* (*U*), *whose density is denoted by p*(*u*). *Intervening on P replaces only its assignment*.

The SCM describes the additive transition and NB decoder at fixed global quantities. It does not introduce a separate generative mechanism for the auxiliary classifier.

##### Definition 4

(Potential outcome of a perturbation). *For a noise realization u, the expression under intervention do*(*P* = *p*) *is*

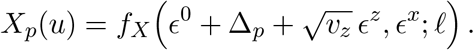

##### Definition 5

(Potential outcome under control). *The null-perturbation outcome is*

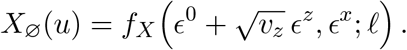

*In particular, control counts are decoded from the control expression latent Z*_∅_, *not directly from Z*^0^.

For integer count vectors *x* and *x*^0^, the joint counterfactual probability is

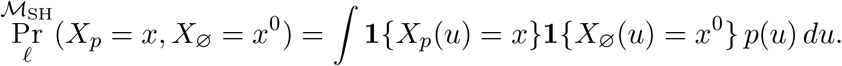

Whenever Pr_*l*_(*X*_∅_ = *x*^0^) *>* 0, abduction gives

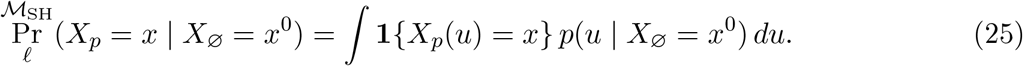

Conditioning on the control counts generally updates all outcome-generating noises. Replacing this posterior by a background posterior times the unconditional priors of *ϵ*^*z*^ and *ϵ*^*x*^ is not valid for this shared-noise counterfactual. Define the control expression latent 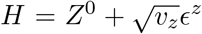, whose distribution is *ν*_0_ = *N*(0, (1 + *v*_*z*_)*I*_*d*_). Under intervention do(*P* = *p*), the expression latent is therefore *Z*_*p*_ = *H* + Δ_*p*_. The counterfactual estimand is given by Eq. (25).

#### 4.6.3 Causal Estimand Identifiability

An estimand is identifiable within a specified class of SCMs if every model in that class inducing the same observed distribution gives it the same value. We state explicit sufficient assumptions here.

##### Assumption 1

(Causal validity and treatment coverage). *The fixed-offset data distribution is generated by an additive SCM of the form in Definition 3, with consistent treatment labels, no interference between cells, and assignment P independent of the outcome-generating exogenous variables within the experimental context. Every perturbation considered, including control, has positive assignment probability. The observed outcome satisfies X* = *X*_*P*_ .

Under this assumption, the marginal interventional distribution is identified by Pr_*l*_(*X*_*p*_ = *x*) = Pr_*l*_(*X* = *x* | *P* = *p*). This fact alone does not identify the joint distribution of potential outcomes for the same cell.

##### Assumption 2

(Identifiability of the noisy observation model). *For any two admissible models inducing the same conditional count distribution for all p* ∈ P, *their observation kernels K*_*l*_ *and K*_*l*_ *are related by one common bijection T* : ℝ^*d*^ → ℝ^*d*^:

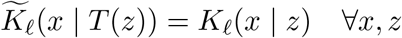

*Moreover, the mixing operator of each kernel is injective on the considered latent distribution with finite first moments. If two such distributions ν*_1_, *ν*_2_ *satisfy*

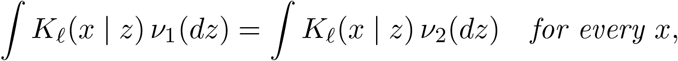

*then ν*_1_ = *ν*_2_. *The class of considered distributions includes their transforms under the admissible coordinate changes*.

##### Assumption 3

(Common affine alignment). *The coordinate change in Assumption 2 can be chosen as T* (*z*) = *Sz* + *c, where S* = ΠΛ, Π *is a permutation matrix*, Λ *is an invertible diagonal matrix, and c* ∈ R^*d*^. *The same transformation applies to the control and every perturbation*.

When both models retain the fixed control distribution (0, (1 + *v*_*z*_)*I*_*d*_) with the same *v*_*z*_, admissible affine changes are further restricted to *c* = 0 and *SS*^T^ = *I*_*d*_, hence to signed permutations within this affine family.

##### Lemma 1

(Identifiability of additive shifts up to aligned coordinates). *Suppose two models satisfy Assumptions 1–3 and induce the same Prl(X* | *P = p) for every p ∈ P. Let νp and* 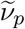 *be their latent expression distributions under p, and let T* (*z*) = *Sz* + *c be the common alignment. Then*

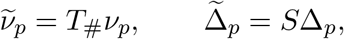

*where T*_#_*νdenotes the pushforward of νunder T*.

*Proof*. Equality of the observed conditional distributions and the kernel relation imply

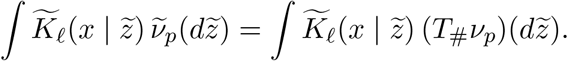

Injectivity of the mixing operator gives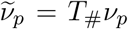, including for control. Additivity gives *Zp* = *H* + Δ_*p*_ and Δ_∅_ = 0. Therefore, writing E_*ν*_*p* [*Z*] for the mean under *ν*_*p*_,

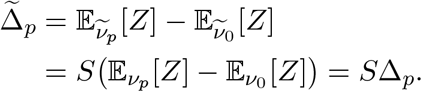

The row covariance of perturbation embeddings is a prior over global effect parameters. Conditional on these parameters, it does not itself introduce cell-level confounding. Absence of assignment confounding follows from Assumption 1. It also does not imply conditional independence of potential outcomes.

##### Assumption 4

(Specified coupling across interventions). *All admissible SCMs reuse the same control expression latent H across interventions, with Z*_*p*_ = *H* + Δ_*p*_, *and use the coordinate-wise inverse-CDF observation mechanism defined above with the same ϵ*^*x*^ *in every potential outcome*.

##### Theorem 1

(Identifiability of the SHERLOCK counterfactual estimand). *Under Assumptions 1, 2, 3, and 4, the joint distribution* Pr_*l*_(*X*_*p*_1, …, *X*_*p*_*k*) *for any finite collection of represented perturbations is identifiable from the fixed-offset observed distribution* Pr_*l*_(*X, P*) *within this restricted model class. Consequently, Eq*. (25) *is identifiable whenever the conditioning control event has positive probability*.

*Proof*. For target count vectors *x*_1_, …, *x*_*k*_, the specified coupling gives

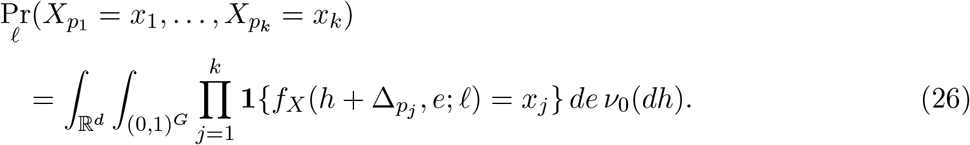

By Lemma 1, any observationally equivalent admissible model has 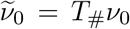and 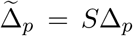. Hence (*h*)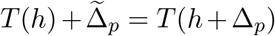. Equality of the product negative bionomial kernels under *T* implies equality of each gene’s conditional CDF. Let ℳ and 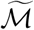 be two admissible SCMs inducing the same observed distribution. The equality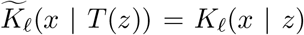 implies equality of the conditional marginal CDFs for every gene:

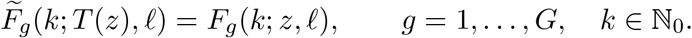

Because both SCMs use the specified inverse-CDF observation mechanism, their outputs agree when evaluated with the observation noise held fixed:

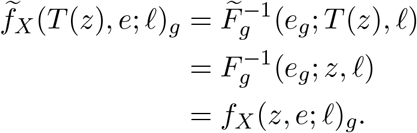

Thus 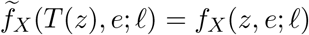for almost every *e* under the uniform distribution on (0, 1) ^*G*^.

By Lemma 1, 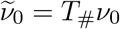 and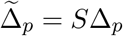. Since *T* (*h*) = *Sh* + *c*,

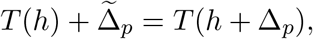

and consequently

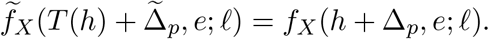

Changing variables via 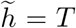 (*h*) in Eq. (26) therefore preserves every outcome indicator and the integrated probability. Including control among the interventions gives

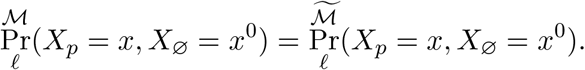

Hence, for any *x*^0^ such that

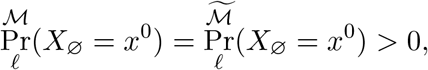

the conditional probabilities satisfy

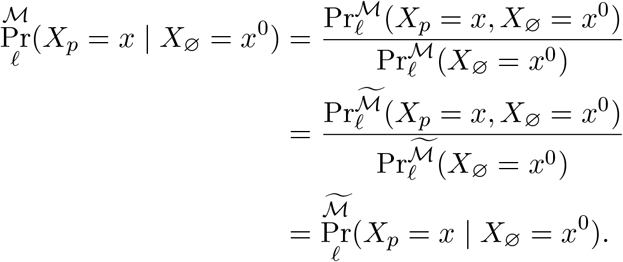

Thus, all observationally equivalent admissible SCMs agree on the conditional counterfactual distribution, establishing identifiability within the specified model class.

The theorem is a sufficient-conditions result, not an unconditional identifiability guarantee for our method. While our framework establishes theoretical identifiability under strict conditions, bridging these guarantees to finite-sample deep learning involves inherent empirical approximations. In practice, enforcing an exact injective mapping is challenging due to finite sample sizes, dropout, and high noise level in single-cell transcriptomics, which can lead to mild local non-identifiabilities or boundary bleeding in the latent space. Similarly, while structural sparsity regularization effectively breaks rotational invariance and prevents arbitrary mixing, achieving exact axis-alignment depends heavily on hyperparameter tuning and the richness of the perturbation dataset. Identifiability experiment results across random initializations and finite samples are provided to support our argument for estimation robustness.(Fig. S1, and S7)

#### 4.6.3 Causal Effect Estimation with SHERLOCK

The implemented causal effect estimation produces deterministic expression-profile predictions. We retain the abduction–intervention–prediction organization while stating the actual computations.

##### Abduction

Let 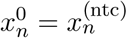be a control profile. Write

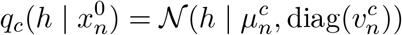

for the variational distribution at the control expression-latent site *z*, where 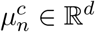and 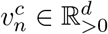 are returned by the control encoder after the prescribed normalization. The guide shares these parameters with the control *z*^0^ site, although their samples are distinct. Thus the helper returning the *z*^0^ encoder mean also returns the control *z* mean. We interpret this output as an approximation to the control expression latent *H*, since control counts are decoded from *z*. The implementation uses only 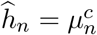; it does not recover the joint posterior of all exogenous noises.

##### Intervention

Let *ρ* be the global perturbation embedding matrix and *q*(*ρ*) its variational distribution. Denote its posterior mean by *m*^*ρ*^ = E_*q*_[*ρ*], the learned row-covariance Cholesky factor by *C*_*P*_, and the deterministic gate output by *W* det. Define *Â* = *C*_*P*_ *m*^*ρ*^ and 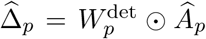.The implemented intervention is

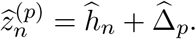

If the full control variational distribution were propagated instead, the shared additive noise construction would give

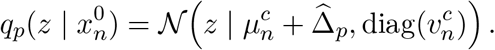

There is no additional independent transition-noise draw here: *H* already includes the factual transition residual, which is held fixed across worlds induced by interventions.

##### Prediction

Holding an offset *l* fixed, we compute

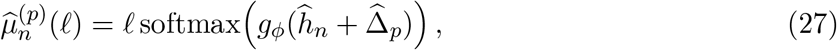

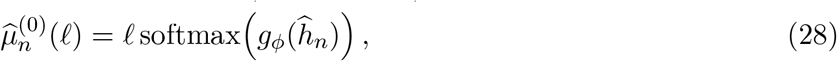

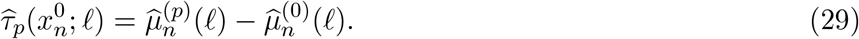

The NB stabilizer cancels in this difference. The decoder outputs logits, so *g*_*ϕ*_(*z*) itself is not an expected count vector.

The corresponding model-defined conditional mean expression difference is

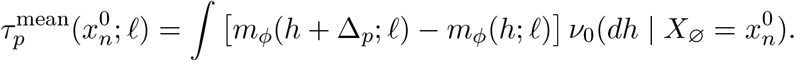

For perturbation *p* at a fixed library-size offset *l*, define the gene-wise average treatment effect (ATE) in the target cell population as

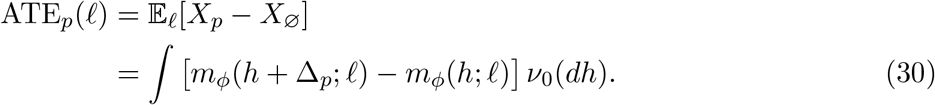

By iterated expectation, averaging the posterior-integrated mean-profile contrast over the population distribution of control profiles recovers this quantity:

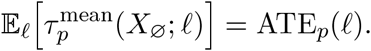

In practice, we approximate the control latent posterior by *q*_*c*_ and evaluate the decoder at its mean. For *N*_0_ control profiles representative of the target population, averaging the resulting predicted contrasts gives the model-based plug-in ATE estimator

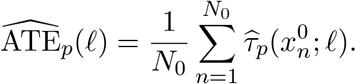

Its accuracy depends on model specification, posterior approximation, and the representativeness of the control profiles. Evaluating the nonlinear decoder at the posterior mean introduces an additional approximation relative to posterior integration. This estimator targets the population-average perturbation effect at the chosen offset; it does not establish recovery of individual-cell counterfactual effects.

### 4.7 Combinatorial perturbations and genetic interaction scoring

The sections above describe the single-perturbation case in which each cell carries exactly one perturbation label *p*_*n*_. We now extend the model to data in which cells may be exposed to a *pair* of perturbations simultaneously, and introduce a learned interaction term that captures non-additive effects.

#### 4.7.1 Notation and input encoding

In the combinatorial setting each cell *n* is associated with an ordered pair 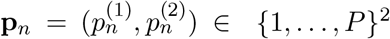, where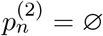 (encoded as −1) for single-perturbation cells. Perturbation labels are indexed at the level of individual genes: a label A+B is split into its components, so a combination introduces no parameters of its own and instead reuses the embeddings, gates and covariance of its two constituents. Let

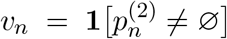

indicate that cell *n* carries a second perturbation. All remaining notation, *A, W*, *ρ, z*^0^, *z, θ*, is unchanged from Sections 4.1–4.4; in particular 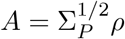 is the covariance-transformed perturbation embedding and *W* ∈ [0, 1] ^*P* ×*d*^ the gate.

#### 4.7.2 Combinatorial latent shift

For a single-perturbation cell the gated shift is unchanged,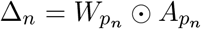. For a combinatorial cell the shift is a reweighted sum of the two parent shifts plus a learned interaction residual

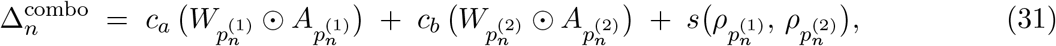

with *c*_*a*_ = *c*_*b*_ = 1 and *s* = 0 recovering the additive baseline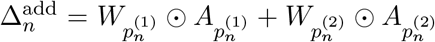 Single-perturbation cells bypass (31) entirely and retain 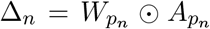. The shift enters the latent transition exactly as in the single-perturbation case, with 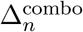substituted for Δ_*n*_; the decoder, KL terms and remaining ELBO structure are unchanged.

##### Interaction residual

The residual *s* is a two-layer perceptron applied to the concatenated pair of perturbation embeddings and symmetrised over both orderings:

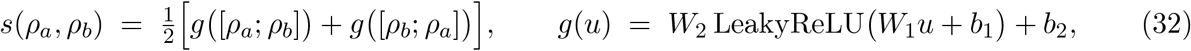

with *W*_1_ ∈ ℝ^*h*×2*d*^, *W*_2_ ∈ ℝ^*d*×*h*^ and hidden width *h*. Averaging the two orderings enforces *s*(*ρ*_*a*_, *ρ*_*b*_) = *s*(*ρ*_*b*_, *ρ*_*a*_), so `A+B` and `B+A` induce identical shifts. The residual is unconstrained in direction and is therefore free to point outside the span of the two parent shifts, which is required to represent asymmetric masking (epistasis) and transcriptional states absent from either single perturbation (neomorphic effects).

##### Parent coefficients

The scalar coefficients *c*_*a*_, *c*_*b*_ rescale each parent’s contribution in the context of the combination. Both are produced by a single bias-free linear map *w* ∈ ℝ^2*d*^ applied to the ordered pair, evaluated in both orderings:

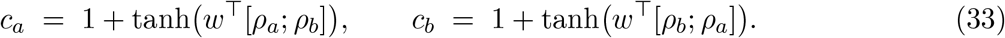

This construction has the two properties the interaction classes require. It is swap-equivariant, *c*_*a*_(*ρ*_*a*_, *ρ*_*b*_) = *c*_*b*_(*ρ*_*b*_, *ρ*_*a*_), giving the label-order invariance appropriate to an unordered pair. And because the map acts on the full ordered pair, the two coefficients vary independently and their sum is unconstrained, so a combination can scale its additive component as a whole—both parents amplified, as in potentiation, or both damped, as in suppression—as well as redistribute weight between them, as in dominance. The tanh confines each coefficient to (0, 2), so a parent’s program may be damped or up to doubled but cannot change sign or diverge, and *w* is initialized near zero so that *c*_*a*_ = *c*_*b*_ ≈ 1 at initialization.

#### 4.7.3 Additivity prior and objective

Nothing in the reconstruction likelihood penalizes non-additivity, so without an explicit prior the model can introduce interaction structure into genuinely additive pairs at negligible cost to the ELBO—which would erase the additive signature that the genetic-interaction scoring below reads out. We therefore add two quadratic penalties over combinatorial cells:

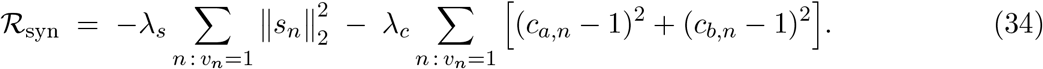

Both accumulate over the cells carrying a combination, as its likelihood contribution does, so how much evidence a pair must show before departing from additivity does not depend on how many cells happen to carry it. Together with the near-zero initialization of *w*, these make the additive model the default: the model begins exactly at Δadd and must be driven away from it by the data. The complete objective is

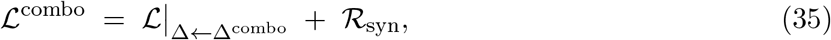

where ℒ is (17) evaluated with the combinatorial shift (31). The auxiliary classifier loss is the one component that changes form: the single-label cross-entropy is replaced by binary cross-entropy against a multi-hot target **t**_*n*_ ∈ {0, 1}^*P*^ with *t*_*n,p*_ = 1 for each perturbation present in cell *n*,

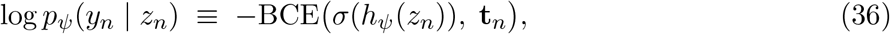

with *σ* the element-wise sigmoid.

#### 4.7.4 Effect vectors for interaction scoring

Interaction scoring operates on effect vectors in gene space—one per single perturbation and one per pair. In the main analysis each vector is taken from the model’s approximate posterior over that perturbation’s own cells: we decode *q*(*z*) for every cell carrying the perturbation, average the decoded profiles, and subtract a non-targeting control (NTC) baseline obtained the same way from the full NTC population. All three vectors entering a comparison, *δ*_*a*_, *δ*_*b*_ and *δ*_*ab*_ ∈ ℝ^*G*^, are referenced to this single shared baseline and expressed as log_2_ CPM differences, so a null perturbation maps to 0 and the additive residual is not offset by a baseline mismatch. Perturbations with fewer than five observed cells are not scored.

The alternative is to generate each effect by applying the corresponding latent shift— (31) for a pair—to a background abducted from NTC cells and decoding, reusing the same background throughout. Both estimators target the marginal interventional means identified by Lemma 2, so the choice between them is one of estimator behavior rather than of what is identified. We score from the posterior for two reasons. Generating all three vectors makes the regression close to degenerate by construction: the double is then a reweighted sum of the singles plus the interaction residual, so the fitted coefficients and residual recover the model’s own parameters rather than a comparison against data. And the reference categories were assigned from measured joint profiles, so estimating the joint effect from the cells that received the pair mirrors the construction those labels came from.

For combinations withheld from training we instead generate the pair’s effect from (31) while retaining posterior-decoded singles, so that the pair is predicted from its two constituents and its own cells are not used. Regressing a generated double against observed singles also keeps the interaction residual discriminative.

The status of this setting should be stated precisely, since it differs from the rest of the analysis. Identification is relative to the distribution a model is fitted to: withholding a combination removes the very conditional that identifies its mechanism, so the resulting prediction is an extrapolation rather than an identified counterfactual, obtained by applying parent coefficients and an interaction residual fitted on other pairs. What makes the extrapolation possible is structural, a combination reuses its constituents’ parameters and therefore has a fully specified shift whether or not it was observed, and whether it is accurate is an empirical question. We make no identification claim for withheld combinations. Because such combinations were nonetheless assayed, their effects are independently measured, and we evaluate the extrapolation against that measurement in two ways: as the correlation between predicted and measured effect profiles, and as the accuracy of the interaction classes assigned from the generated joint effect.

#### 4.7.5 Genetic interaction metrics

We quantify the interaction type of each pair by Norman-style regression in decoded gene-expression space. Writing the additive prediction as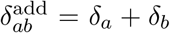, the double effect is regressed on its two single effects,

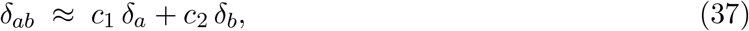

by Theil–Sen regression without intercept, giving coefficients (*c*_1_, *c*_2_) and fitted double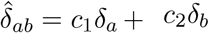. Note that (*c*_1_, *c*_2_) are post hoc regression coefficients in gene space and are distinct from the model’s parent coefficients (*c*_*a*_, *c*_*b*_) of (33). Table 1 lists the eleven scalar metrics derived from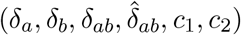. The same routine is applied to the effect vectors of any method under comparison, so that only the effect vectors, and not the scoring convention, differ between methods.

**Table 1.**
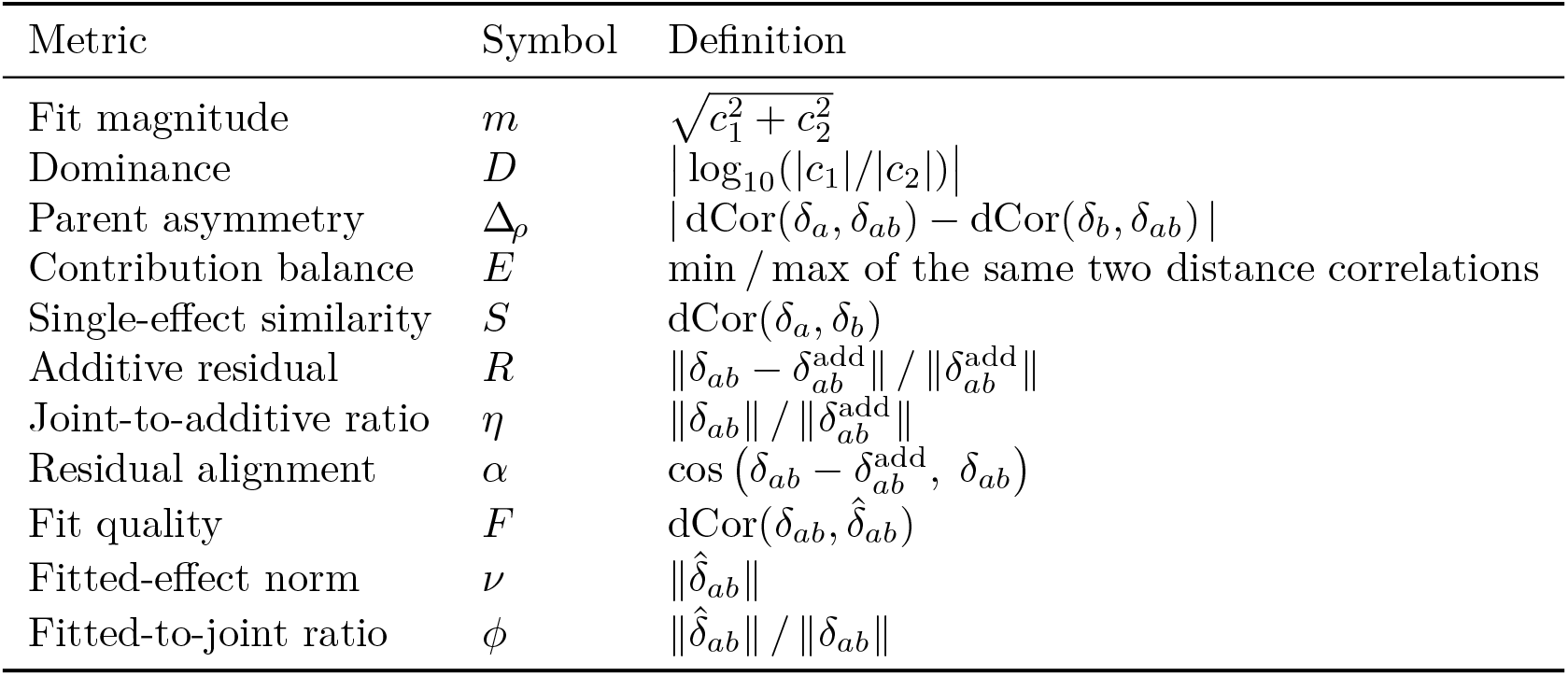
Genetic-interaction metrics. dCor denotes distance correlation.

#### 4.7.6 Interaction classification

Each metric is *z*-scored across the pairs being classified. Every class is assigned a composite score equal to the mean of its signed *z*-scored terms (Table 2), and each pair receives the class with the highest score. Pairs with any non-finite metric are left unclassified. The reference annotation distinguishes two synergy subtypes by whether the two single perturbations induce similar or dissimilar phenotypes; we do not make this distinction and score synergy as one class throughout, collapsing the reference subtypes accordingly.

**Table 2.** Composite scores over *z*-scored metrics; each is averaged over its terms.

| Class | Biological interpretation | Composite score |
| --- | --- | --- |
| Approximately additive | Double effect explained by the sum of the singles | $-R - \eta$ |
| Epistasis | One parent dominates and masks the other | $+D + \Delta_\rho - E$ |
| Redundant | Parents act on a shared program; the double adds little beyond either alone | $+S - \eta$ |
| Suppression | Joint response damped relative to what both parents predict | $-\eta + \phi - m$ |
| Neomorphic | Double not explained by any weighting of the parents | $-F - \nu$ |
| Potentiation | Double exceeds the additive expectation; parents act on dissimilar programs | $+\eta + \alpha - S$ |
| Synergy | Double exceeds the additive expectation; parents act on similar programs | $+m + \eta + 2S$ |

#### 4.7.7 Theoretical extension to combinatorial perturbations

The Theorem above is stated for an arbitrary set of target perturbations. To apply it to combinations we first make explicit that a combination is an intervention in ℳ_*SH*_, and that the quantification in Lemma 1 ranges over the perturbations the assay actually contains.

##### Definition 6

(Combinatorial interventions and observed support). *Let G denote the set of individual perturbation targets. We take the perturbation variable to be the pair P* = (*P*_1_, *P*_2_), *with P*_2_ = ∅ *for singly-perturbed cells, so that the latent mechanism reads Z* = *f*_*p*_(*Z*^0^, *P*_1_, *P*_2_, *ϵ*^*z*^). *A combination of a, b* ∈ *G is the joint intervention do*(*P*_1_ = *a, P*_2_ = *b*), *which is admissible under Definition 2 since that definition applies to any X* ⊆ *V* ; *we abbreviate it do*(*P* = *ab*) *and write f*_*ab*_(·) := *f*_*p*_(·, *a, b*, ·) *for the induced mechanism. We write*

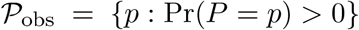

*for the set of perturbations, single or combinatorial, appearing in the data, and take the quantifica-tion* ∀p_*i*_ ∈ P *in Lemma 1 to range over* P_obs_.

##### Lemma 2

(Identifiability of combinatorial interaction estimands). *Let a, b* ∈ *G and suppose Assumptions 2 and 3 hold with a, b, ab* ∈ *P*_obs_. *Then*

1. *the joint counterfactual distribution p*^*ℳ Sℋ*^ (*X*_*a*_, *X*_*b*_, *X*_*ab*_) *is uniquely identifiable from the observational distribution p*(*X, P*);
2. *the marginal interventional means µ*_*q*_ = E[*X*_*q*_] *for q* ∈ {*a, b, ab*, ∅} *are identifiable; and*
3. *every genetic-interaction metric computed from the effect vectors δ*_*q*_ = *µ*_*q*_ − *µ*_∅_ *is identifiable*.

*Proof*. (i) By Definition 6, *ab* is an intervention in ℳ_*SH*_, and *a, b, ab P*_obs_, so the hypothesis of Lemma 1 is satisfied for each: the mechanisms *f*_*a*_, *f*_*b*_, *f*_*ab*_ and the inverse decoder (*f*^*X*^) ^™1^ are identified up to the common permutation and scaling ΠΛ fixed by the Lemma. The Theorem then applies verbatim with ℙ = {*a, b, ab*} and *k* = 3.

(ii) Marginals of an identified joint distribution are identified, so each *p*(*X*_*q*_) and hence each *µ*_*q*_ = E[*X*_*q*_] is identified.

(iii) Each *δ*_*q*_ is a difference of identified means, and the Theil–Sen coefficients and every derived metric are fixed measurable functions of (*δ*_*a*_, *δ*_*b*_, *δ*_*ab*_); identification is preserved under measurable transformation. Since these quantities are defined in gene space, they are moreover invariant to the residual (Π, Λ) ambiguity of Assumption 3.

##### Remark 1

(Estimation). *Part (ii) concerns* marginal *interventional means, so any consistent estimator of those means identifies the metrics of part (iii). Under randomized perturbation assignment*, E[*X P* = *q*] E[*X P* = ∅] *is such an estimator, and it is the one used in our main analysis. The shared-background construction of the Theorem is required only for per-cell counterfactual contrasts*—t*he distribution of individual treatment effects, or the dependence between potential outcomes within a cell*—w*hich the interaction metrics do not use*.

##### Remark 2

(Non-identifiability of the interaction decomposition). *Lemma 2 identifies the total mechanism f*_*ab*_, *not its decomposition into parent scaling and interaction residual. The residual is unconstrained in direction, so the map* (*c*_*a*_, *c*_*b*_, *s*) ↦ Δ_*ab*_ *is not injective and the three components are not separately identified; the quadratic penalties of the training objective select among observationally equivalent decompositions. All reported quantities depend on* Δ_*ab*_ *alone, and we do not interpret c*_*a*_, *c*_*b*_ *or s individually*.

Lemma 2 requires *ab∈ P*_obs_. For a combination outside the observed support the hypothesis of Lemma 1 is vacuous at *ab*: no observational conditional constrains *f*_*ab*_, and two models agreeing on all observables may disagree there. Identification therefore cannot be recovered, and prediction for such a combination requires an additional assumption, which we state rather than derive.

##### Remark 3

(Compositional transfer). *Let* Δ_*ab*_ = *c*_*a*_(*ρ*_*a*_, *ρ*_*b*_)Δ_*a*_ + *c*_*b*_(*ρ*_*a*_, *ρ*_*b*_)Δ_*b*_ + *s*(*ρ*_*a*_, *ρ*_*b*_) *be the combinatorial mechanism, in which the maps c and s are shared across all pairs. A prediction for ab* ∉ *P*_obs_ *amounts to taking the maps fitted on P*_obs_ *to describe the mechanism of pairs outside it, so that* Δ_*ab*_ *evaluated at* (*ρ*_*a*_, *ρ*_*b*_) *is the mechanism of ab for any a, b* . *It is not implied by Assumptions 2 and 3, and therefore we make no identification claim for combinations outside* _obs_. *It is instead an empirical property of the fitted model, and one that can be measured*.

### 4.8 Interpreting the fitted Norman models

All analyses in this section use the five trained runs described above, the same gene filter (3,270 genes), and Norman’s expert interaction labels with the two synergy subtypes merged (86 of the 88 labeled pairs have all quantities defined). Perturbation-target genes are excluded from every transcriptional readout, so a CRISPRa target can never score its own program. Unless stated otherwise, a quantity reported for a pair is the mean of its five per-run values.

#### 4.8.1 Latent factors: gates, decoded signatures and matching across runs

For each run we take the deterministic gate *W* ∈ [0, 1] ^*P*×*d*^ (*d* = 16) and the gated shift Δ_*p*_ = *W*_*p*_ ⨀ *A*_*p*_ of every perturbation (Section 4.7), evaluated with the Concrete draw replaced by its deterministic counterpart. A latent factor *j* is *used* by perturbation *p* when *W*_*pj*_ *>* 0.5; factors no perturbation uses are dropped (three of sixteen: F1, F8, F14).

Latent coordinates are exchangeable across runs, so factors are identified by what they do to expression rather than by their index. Writing *g*(·) for the decoder followed by log_2_-CPM normalization and *u* for the latent states abducted from 256 randomly chosen control cells, the signature of factor *j* is the central difference

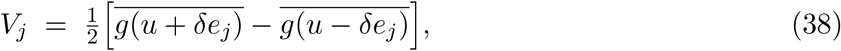

averaged over the 256 backgrounds, with *δ* the median Δ_*pj*_ over the gates that run leaves active and *e*_*j*_ the *j*-th unit vector. Signatures are restricted to readout genes and *l*_2_-normalised. Run 1 is the reference: for each other run we compute the matrix of cos between its signatures and run 1’s, match factors by the Hungarian algorithm on cos, and flip the sign of a matched factor when the cosine is negative. Each run’s gates and shifts are permuted accordingly, so *W* and Δ can be averaged over runs.

Matching quality is reported in **Fig. S14**. For every factor we give the cos of its match in each run, the best alternative partner in the same run, and the distribution of cos between unmatched factor pairs (median 0.09, 95th percentile 0.41), and we test each run’s total matched cos against 5,000 random one-to-one pairings of the same matrix. Two factors (F3, F15) whose best alternative is as similar as their assigned partner are not interpreted.

A factor’s program content is the mean of its signature over a program’s genes, divided by the standard deviation of that signature over all readout genes, so scores are in units of the factor’s own spread. Program gene sets are not taken from a published signature database: they are standard markers of the three lineages K562 can be driven towards, compiled from the haematopoietic differentiation literature [54, 55, 47, 56] and matched to the erythroid, CD41+ megakaryocytic and granulocytic phenotypes Norman et al. report for this screen [43]; every gene used is listed here. On latent factors the erythroid program uses the 20 canonical erythroid genes the model retains (*HBG1, HBG2, HBZ, HBA1, HBE1, HBD, HBQ1, GYPA, GYPB, GYPC, ALAS2, HMBS, UROD, BLVRB, SLC25A37, GATA1, NFE2, TFRC, ANK1, PRDX2*), and the granulocytic program the four canonical granulocyte markers above the gene filter (*CD33, LST1, FCER1G, CST3*). Canonical megakaryocytic markers are all below the filter, so no megakaryocytic score is defined on latent factors. A factor’s loadings are the run-averaged Δ_*pj*_ over perturbations, and a module’s loading is the mean over its members.

#### 4.8.2 Groupings: modules, Norman’s clusters, fate scores and poles

##### Consensus modules

Each run yields a 105 × 105 perturbation correlation matrix. Each run is clustered on 1 − *ρ* by Ward linkage at every resolution *k* = 5, …, 20, and the co-assignment matrix *C*_*ij*_ is the fraction of (run, *k*) pairs in which perturbations *i* and *j* share a cluster. Modules are obtained by average linkage on 1 − *C*. The number of modules is chosen by the largest gain of the consensus over the individual runs, i.e. the *k* maximising the ratio of the mean ARI between the consensus clustering and the per-run clusterings to the mean run-to-run ARI, giving 12 modules, numbered by decreasing size.

##### Norman’s clusters and phenotype groups

Perturbation-map clusters are taken from Table S6B of [43]; clusters with at least two members in the screen are kept. Three named phenotype groups are added from the same study: erythroid (*CBL, CNN1, UBASH3A, UBASH3B, PTPN9, PTPN12*), granulocyte (*CEBPA, CEBPB, CEBPE, SPI1*) and G1 arrest (*CDKN1A, CDKN1B, CDKN1C*), giving 20 groups in total. For each group we record the share of its members falling in the single module that holds most of them, and compare it with the same statistic under 5,000 random permutations of the module labels across perturbations, which preserves module sizes and group sizes. The 20 *p*-values are adjusted together by Benjamini-Hochberg, and each group’s share and *q*-value are shown with the cluster-by-module overlap (**Fig. S12a**).

##### Observed fate scores

Fate scores are computed from the measured counts before the model’s gene filter, so that markers below the filter still contribute. Marker sets are the same literature-derived lineage panels used above [54, 55, 47, 56, 43], restricted to genes that are not themselves CRISPRa targets: erythroid (globins, glycophorins, *ALAS2, SLC25A37, GATA1, EPOR, AHSP, SLC4A1* ; 15 genes), megakaryocytic (*ITGA2B, ITGB3, GP9, GP1BA, PF4, PPBP, VWF, TUBB1, MPL, CD36, PECAM1, PLEK* ; 12 genes) and granulocytic (*MPO, ELANE, AZU1, PRTN3, CTSG, LYZ, S100A8/9, CSF3R, ITGAM, CD33, MNDA, LST1, FCER1G, CST3, LGALS3, ANXA1* ; 16 genes). For every perturbation and marker we take log_2_ of pseudobulk CPM plus 0.05 minus the same quantity in control cells, *z*-score each marker across the 105 perturbations, and average over a lineage’s markers. The model’s decoded erythroid score is the same program average applied to the decoded single-perturbation effects.

Because megakaryocytic markers are detected in only about 1% of control cells, the reliability of each score is measured directly: cells of every perturbation and of the controls are split into two random halves, scores are recomputed within each half, and the two halves are correlated across perturbations (10 random splits), with the Spearman-Brown correction [57] giving the full-data reliability. The correlation between the erythroid and megakaryocytic scores is reported with a bootstrap confidence interval over perturbations (5,000 resamples), a permutation *p*-value (5,000 shuffles of one score) and a recomputation with the strongest megakaryocytic perturbation removed.

##### Poles and identities

A module’s fate scores are the means over its members. Modules with a mean erythroid score above +0.2 form the *erythroid-high* pole, those below 0.2 the *erythroid-low* pole, and the rest are neutral. Within the down pole, a module is called granulocytic when its granulocytic score exceeds 0.5, and erythroid-low otherwise; within the up pole, a module is called erythroid plus megakaryocytic when its megakaryocytic score exceeds 0.3. The cut-offs are round numbers chosen for readability, not fitted.

##### Interaction enrichment

A labeled pair is described by the poles of its two parents’ modules, by whether both parents belong to the same module, and by whether the parents’ observed erythroid scores have opposite signs. Each claim in the Results compares one interaction class against all other labeled pairs by Fisher’s exact test, and the eight tests are adjusted together by Benjamini–Hochberg. These tests describe the measured, labeled panel, which is a curated selection of pairs; they are not extrapolated to unmeasured combinations.

#### 4.8.3 Reading interaction type from the model’s parameters

##### Descriptors

Let *a* and *b* index the parents of a labeled pair, Δ_*a*_, Δ_*b*_ their gated shifts, *c*_*a*_, *c*_*b*_ their parent coefficients, *s* the interaction residual, and 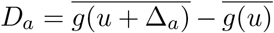 the decoded single-perturbation effect (256 control backgrounds, as above). Write Δ_*ab*_ = *c*_*a*_Δ_*a*_ + *c*_*b*_Δ_*b*_ + *s* for the combined shift, *D*_*ab*_ for its decoded effect, and *m*_*a*_ = **1**[*W*_*a*_ *>* 0.5] for the factors a parent uses. The seven descriptors are

- **Direction** (rho), *ρ* = entry (*a, b*) of the run-averaged perturbation correlation matrix: do the two perturbations push the cell the same way?
- **Private factors** (`private_frac`), 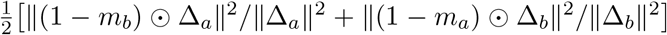 : how much of each parent’s effect runs through latent factors its partner does not use?
- **Parent scales** (`c_sum`), *c*_*a*_ + *c*_*b*_: does the model keep, damp or amplify each parent’s program in the pair?
- **Interaction size** (`syn_rel`), ‖*s*‖*/* ‖Δ_*a*_ + Δ_*b*_‖: how large is the non-additive term relative to the additive shift?
- **Joint-to-additive ratio** (`joint_over_add`), ‖*D*_*ab*_‖ */* ‖*D*_*a*_ + *D*_*b*_‖: is the decoded combination smaller or larger than the sum of its parents?
- **Dominance** (`dom_gap`), |cos(*D*_*ab*_, *D*_*a*_) −cos(*D*_*ab*_, *D*_*b*_) |: does the combination resemble one parent much more than the other?
- **Off-gate interaction** (`syn_offgate_dec`), 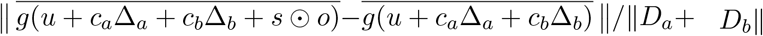: how much of the non-additive term acts on latent factors that neither parent uses?

where *o* = 1 − max(*m*_*a*_, *m*_*b*_) selects the factors neither parent uses. The first three read the perturbation covariance, the gates and the parent-coefficient map; the next two the interaction residual; the last two the decoder. The set was reduced to one descriptor per mechanistic idea—direction, gate sharing, parent scaling, interaction size, size of the combination, dominance, and where the interaction acts—and each is the mean over the five runs. We apply these to the following genetic interactions:

- **Approximately additive**: neither parent is amplified and the interaction term is small.
- **Redundant**: the parents act in the same direction, and the pair does less than their sum.
- **Synergy**: the parents act in the same direction, and the pair does more than their sum.
- **Potentiation**: the parents work through largely separate latent factors and neither is damped, so both programs are expressed in full.

- **Epistasis**: the pair does no more than the sum and looks mostly like one parent.
- **Suppression**: the parents are unrelated or opposed, and the pair does no more than the sum.
- **Neomorphic**: the parents are not aligned, and the interaction term acts on latent factors that neither parent uses.

##### Support for the neomorphic descriptor

Two properties of the fitted model motivate reading neomorphism from the interaction residual. Let 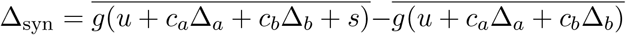 be what the residual adds once decoded. Two quantities are quoted in the Results: the fraction of ‖Δ_syn_‖ lying outside the least-squares span of *D*_*a*_ and *D*_*b*_ (class means 0.59–0.69), and the share of ‖Δ_syn_‖ produced by the off-gate factors *o* (median 0.33), whose Spearman correlations with the gate and direction descriptors are all at most 0.41 in absolute value. Both are computed over all 5,460 single-gene pairs the model can compose, since they describe the fitted model and use no interaction labels; every classification result uses only the 86 labeled pairs.

##### Decision list

Rules are mined as an ordered decision list, because that is how they are applied. Candidate conditions are single thresholds on one descriptor in either direction, taken at the 15th to 85th percentile of that descriptor in 5% steps; percentiles are evaluated over all 5,460 composable pairs so that the grid does not depend on the labels. At each step, and for each class not yet placed, we search all conjunctions of at most two conditions on the pairs no earlier rule has claimed, requiring at least three claimed pairs, and keep the conjunction with the highest *F*_1_ for that class. The class whose best rule has the highest precision is appended to the list, its pairs are removed, and the search repeats until every class has a rule. Pairs no rule claims are called approximately additive, so that class needs no positive signature. The neomorphic rule is additionally required to contain one condition on the off-gate interaction descriptor, in the direction of more novelty; imposing this constraint left overall accuracy unchanged (0.767) and raised neomorphic recall from 0.42 to 0.50, so the mechanistic reading costs nothing.

The mined list is given in Table 3 and is applied with these fixed thresholds; the resulting confusion matrix is **Fig. S11a** (left). Because both the thresholds and the order were selected on the same 86 labeled pairs, the accuracy and balanced accuracy we report measure how well these seven parameters separate the expert classes, and are not estimates of performance on unseen pairs. Per-class separability is additionally summarized by the one-versus-rest AUROC of each descriptor, computed as the Mann-Whitney *U* statistic divided by the product of the class sizes (**Fig. S11a**, right), and each class’s mean descriptor profile is shown in **Fig. S11b**.

**Table 3:** Decision list mined on the 86 labeled Norman pairs. Rules are applied top to bottom; each is evaluated only on the pairs no earlier rule has claimed, and the pairs no rule claims are called approximately additive. Precision and the number of pairs claimed are measured at the point in the list where the rule is applied. Overall accuracy 0.767, balanced accuracy 0.781.

| Class | Rule (applied in this order) | Precision | Pairs claimed |
| --- | --- | --- | --- |
| approximately additive | $c\_sum \leq 1.877$ and $syn\_rel \leq 0.173$ | 0.88 | 8 |
| redundant | $\rho > 0.360$ and $joint\_over\_add \leq 0.966$ | 0.86 | 7 |
| synergy | $\rho > 0.360$ and $joint\_over\_add > 1.101$ | 0.83 | 23 |
| potentiation | $private\_frac > 0.249$ and $c\_sum > 1.503$ | 0.86 | 7 |
| epistasis | $joint\_over\_add \leq 1.031$ and $dom\_gap > 0.643$ | 0.70 | 10 |
| suppression | $\rho \leq 0.135$ and $joint\_over\_add \leq 1.136$ | 0.79 | 14 |
| neomorphic | $syn\_offgate\_dec > 0.065$ and $\rho \leq 0.326$ | 0.75 | 8 |

### 4.9 Datasets and Preprocessing

#### 4.9.1 Sciplex EGFR drug perturbation data analysis

BT333 glioblastoma cells treated with EGFR/kinase-pathway inhibitors or DMSO at 10 *µ*M were restricted to perturbations with *>* 100 cells and to genes differentially expressed between each drug and DMSO (Wilcoxon test, FDR *<* 0.05), yielding 11,420 cells and 1,157 genes. Perturbation effects were defined as library-size-normalized, log-transformed expression shifts relative to DMSO. SHERLOCK was fit to these data with a latent dimension of 16, batch size of 4,096, L0 sparsity penalty *λ* = 50, and a maximum of 1,000 epochs with early-stopping patience of 350 epochs.

#### 4.9.2 Replogle dataset analysis

We used the genome-scale CRISPRi Perturb-seq screen of K562 cells from Replogle et al. [20] (10x Chromium). The working subset comprises 118,641 cells and 1,185 genes over 682 perturbation labels (681 targeted genes plus a non-targeting control), with the non-targeting label mapped to the model’s control label and no further cell or gene filtering. As an external ground truth for perturbation grouping we used the eight non-overlapping complexes and pathways annotated in the original study, covering 335 targeted genes: exosome (20 genes), Mediator complex (26), mitochondrial protein translocation (40), nucleotide excision repair (23), 39S (43), 40S (97) and 60S (53) ribosomal subunits, and spliceosome (33). No perturbation belongs to more than one group. Per-perturbation effects were computed as the difference in mean log_2_-CPM expression between each perturbation’s cells and the non-targeting cells.

SHERLOCK was fit with a 16-dimensional latent space, batch size 4096, hard-concrete (L0) gate penalty *λ* = 0.1 ramped in over the first 300 epochs, and an auxiliary perturbation-classification weight of 20, for up to 200 epochs (500 in the reproducibility analysis) with validation every 50 epochs and patience 20. Five independently seeded runs were trained; the baselines (cVAE, sVAE+, contrastiveVI, scGen) used the same latent dimension, batch size and number of runs.

#### 4.9.3 RAEFISH dataset analysis

Imaging-based spatial RNA profiling (RAEFISH) [23] provided 17,931 segmented cells assayed over a 492-gene panel, spanning five imaging experiments and 551 perturbation labels, with per-cell barcode-calling fields (identity and read counts of the best and second-best barcode), MERFISH and RAEFISH totals and spatial coordinates. Because most perturbations are present in only a subset of experiments (93 of 551 occur in a single experiment), analysis was restricted to the single largest experiment (5,342 cells) so that perturbation effects are not confounded by batch. Within it we kept perturbations with at least 20 cells (3,898 cells, 77 labels including the non-targeting control) and then cells with log(total counts + 1) *>* 3, leaving 3,436 cells. All 492 panel genes were retained, and per-perturbation effects were computed as log_2_-CPM differences from the non-targeting cells for the 76 remaining perturbations.

SHERLOCK was fit with a 16-dimensional latent space, a rank-6 shared perturbation covariance, hard-concrete gate penalty *λ* = 10, a linear latent shift, gate temperature annealed from 0.5 to 0.1, batch size 4096 and up to 1,000 epochs.

#### 4.9.4 Sciplex drug perturbation data analysis

Preprocessing. Combinatorial single-cell profiles of BT333 glioblastoma cells (11 drugs + DMSO, ± T-cell co-culture) were quality-filtered by hash-barcode confidence (hash UMI *>* 1, top/second-best ratio *>* 2) and UMI count (*>* 500 per cell). A gene panel was defined per concentration as the union of (i) per-drug DEGs vs. DMSO in T-cell-naive cells (Wilcoxon rank-sum, FDR *<* 0.05) and (ii) the top 1,000 genes most altered by T-cell co-culture within DMSO-treated cells (Wilcoxon, ranked by —score—), yielding a focused gene set (2,894 genes, 35,511 cells at 10 µM, and 1,320 genes, 38,755 cells at 1 µM) for downstream modeling.

Model. We fit Sherlock through a low-rank (rank = 3) shared covariance structure and a hard-concrete (L0) sparse gate over an 8-dimensional latent space, to a perturbation-specific shift applied to a condition-matched control latent background. Models were trained per T-cell condition with batch size 4096; 1,000 epochs; and patience 500.

Perturbation grouping. Pairwise perturbation similarity was defined from the model’s learned latent covariance (*ρ*); perturbations were hierarchically clustered (Ward linkage, 1–*ρ* distance) and grouped at a fixed dendrogram cut, applied consistently across correlation-heatmap and bipartite-graph visualizations.

#### 4.9.5 Sciplex CRISPRi/a perturbation data analysis

Preprocessing. Single-cell CRISPRi and CRISPRa screens comprising ~140 kinase-targeting guides and non-targeting controls (NTCs) were profiled across three T-cell co-culture dose ratios and an untreated condition. Cells were quality-controlled based on sgRNA assignment (*>* 3 sgRNA reads and top-guide proportion *>* 0.3) and hash-barcode confidence, and guides represented by fewer than 50 cells were excluded. Guide-by-treatment metacells were generated by aggregating raw counts from randomly sampled cells without replacement (4 cells per metacell for CRISPRi and 6 cells per metacell for CRISPRa; 30 draws per group). Guides were retained if they yielded *>* 50 significantly differentially expressed genes (DEGs) relative to NTCs in untreated cells (Wilcoxon test, FDR *<* 0.05), resulting in 71 guides plus NTCs. The feature space was defined as the union of the top 50 DEGs for each retained guide and the kinase target genes, yielding 570 genes for CRISPRi and 455 genes for CRISPRa.

Modeling. After excluding untreated metacells, each screen comprised 6,480 metacells (71 guides + NTC × 3 T-cell dose ratios × 30 draws). SHERLOCK was trained separately on the CRISPRi and CRISPRa screens using a latent dimension of 16, covariance rank of 6, *L*_0_ regular-ization coefficient of 35, a maximum of 2,500 epochs, and an early-stopping patience of 500 epochs. Condition conditioning was disabled (`use_conditions=False`), such that metacells from all three T-cell dose ratios were modeled jointly to learn a single set of perturbation effects under T-cell co-culture.

Reproducibility. Each screen was independently trained with five random seeds (N RUNS=5). Reproducibility across runs was assessed using perturbation-clustering co-membership and correlations between estimated counterfactual effect sizes.

#### 4.9.6 Norman dataset analysis

The combinatorial CRISPRa screen of Norman et al. [43] comprises 111,255 K562 cells and 19,018 measured genes, covering 105 single and 131 double gene activations together with non-targeting controls. Genes were retained if their mean expression exceeded 0.5 UMI per cell or if they were themselves a perturbation target, leaving 3,270 genes (3,190 by expression plus 80 target genes recovered by the second criterion), so that every activated gene can be read out even when lowly expressed. Per-perturbation effects were computed as log_2_-CPM differences from the non-targeting cells, giving a 236 × 3,270 effect matrix over the 236 perturbation labels. Genetic-interaction annotations for 88 of the double perturbations were taken from the original study and used only for evaluation.

To test prediction of unseen combinations, 20% of the annotated double perturbations were held out, stratified by interaction class (22 of 88 pairs). All cells carrying a held-out combination were removed from training, leaving 104,626 cells, while every constituent single perturbation was retained, so a held-out pair is predicted entirely from its two parents.

Megakaryocytic priming was a weak signal: its markers are detected in about 1% of control cells, and scoring two independent halves of each perturbation’s cells gave estimates that agreed less well than for the other lineages (0.84, against 0.99 erythroid and 0.97 granulocytic). It still co-varied weakly with the erythroid score(*r* = +0.29 ± 0.11, *q* = 0.009), well below what this level of measurement noise would permit. This is consistent with the shared megakaryocyte-erythroid origin of the two lineages [54, 55].

SHERLOCK was fit in combinatorial mode with the interaction term enabled: a 16-dimensional latent space, a rank-16 shared perturbation covariance, a rank-6 interaction term, hard-concrete gate penalty *λ* = 0.1 ramped in over the first 300 epochs, up to 500 epochs with validation every 50 epochs and patience 30. Five independently seeded runs were trained, and all reported statistics are summarized over those runs. The generative baselines were tuned with Optuna [58], SHERLOCK by grid search, and GEARS with its published Norman settings.

## 5 Acknowledgments

We thank Nick Hou for being the first user of the tool, and Neeha Kothapalli and Aaron Zweig for their helpful discussions and feedback. We also thank Yubao Cheng for sharing the REAFISH dataset and for providing helpful explanations regarding the dataset.

## 7 Author Contributions

M.Z., J.D.M., L.S., J.L.M., and E.A. conceived the study, and E.A. and J.L.M. provided overall supervision of the study. M.Z., J.D.M., L.S., J.L.M., and E.A. designed and developed the SHERLOCK framework. M.Z., J.D.M., L.S., E.A., and J.L.M. contributed to implementation. M.Z., J.D.M., L.S., J.L.M., and E.A. analyzed data. M.Z., J.D.M., L.S., and E.A. contributed to the theoretical analysis. J.L.M., L.S., and R.M.G. contributed to the EGFR inhibitor screen and its analysis. M.Z., J.D.M., S.C. contributed to benchmarking. M.Z., J.D.M., L.S., R.M.G., S.C., J.L.M., and E.A. interpreted data. M.Z., J.D.M., L.S., J.L.M., and E.A. wrote the paper. All authors reviewed, contributed to, and approved the paper.

## 8 Lead Contact

Further information and requests for resources should be directed to and will be fulfilled by the Lead Contacts, Jose L. McFaline-Figueroa, and Elham Azizi.

## 9 Data Availability

The Norman et al. Perturb-seq dataset analyzed in this study is publicly available through Figshare (https://figshare.com/articles/dataset/Norman_et_al_2019_Science_labeled_Perturb-seq_data/24688110?file=43390776). The Replogle et al. Perturb-seq dataset is publicly available through the Genome-Wide Perturb-seq portal (https://gwps.wi.mit.edu). The spatial perturbation dataset is publicly available through Mendeley Data (https://data.mendeley.com/datasets/8kbv637pxh/1). The EGFR inhibitor screening dataset is publicly available through the Gene Expression Omnibus (GEO) under accession number GSE261618. The GBM kinase genetic and drug screening datasets will be deposited in GEO and made publicly available upon publication.

## 10 Code Availability

The source code for SHERLOCK and associated documentation are available at https://github.com/azizilab/SHERLOCK. Code and scripts used to reproduce the analyses and figures presented in this study are available at https://github.com/azizilab/SHERLOCK_Reproducibility.

## Supplementary Figures

**Figure S1:**
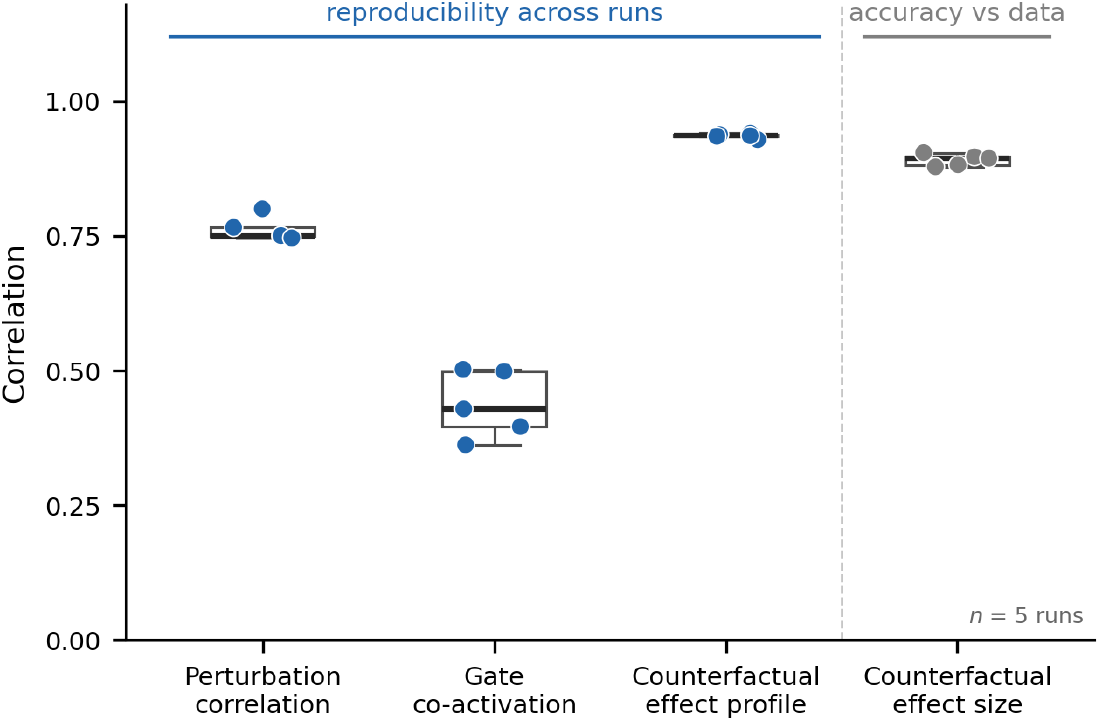
Evaluating reproducibility across five independently trained runs (Replogle screen[20]). Each point is one run; the box shows the spread of the five values. The first three columns measure reproducibility across runs: each run is compared with the consensus of the other four (leave-one-out, Spearman). The fourth measures accuracy against the data within each run (Pearson). *Perturbation correlation*: the 681±681 matrix of learned similarities between perturbations (0.763±0.022). *Gate co-activation*: for every pair of perturbations, how much their sets of gated latent factors overlap; this is compared instead of the gates themselves because latent coordinates are exchangeable between runs (0.438 ± 0.062). *Counter-factual effect profile*: the predicted perturbation-by-gene effect matrix (0.936 ± 0.004). *Counterfactual effect size*: the predicted per-perturbation effect magnitudes against the measured ones (0.891 ± 0.011).

**Figure S2:**
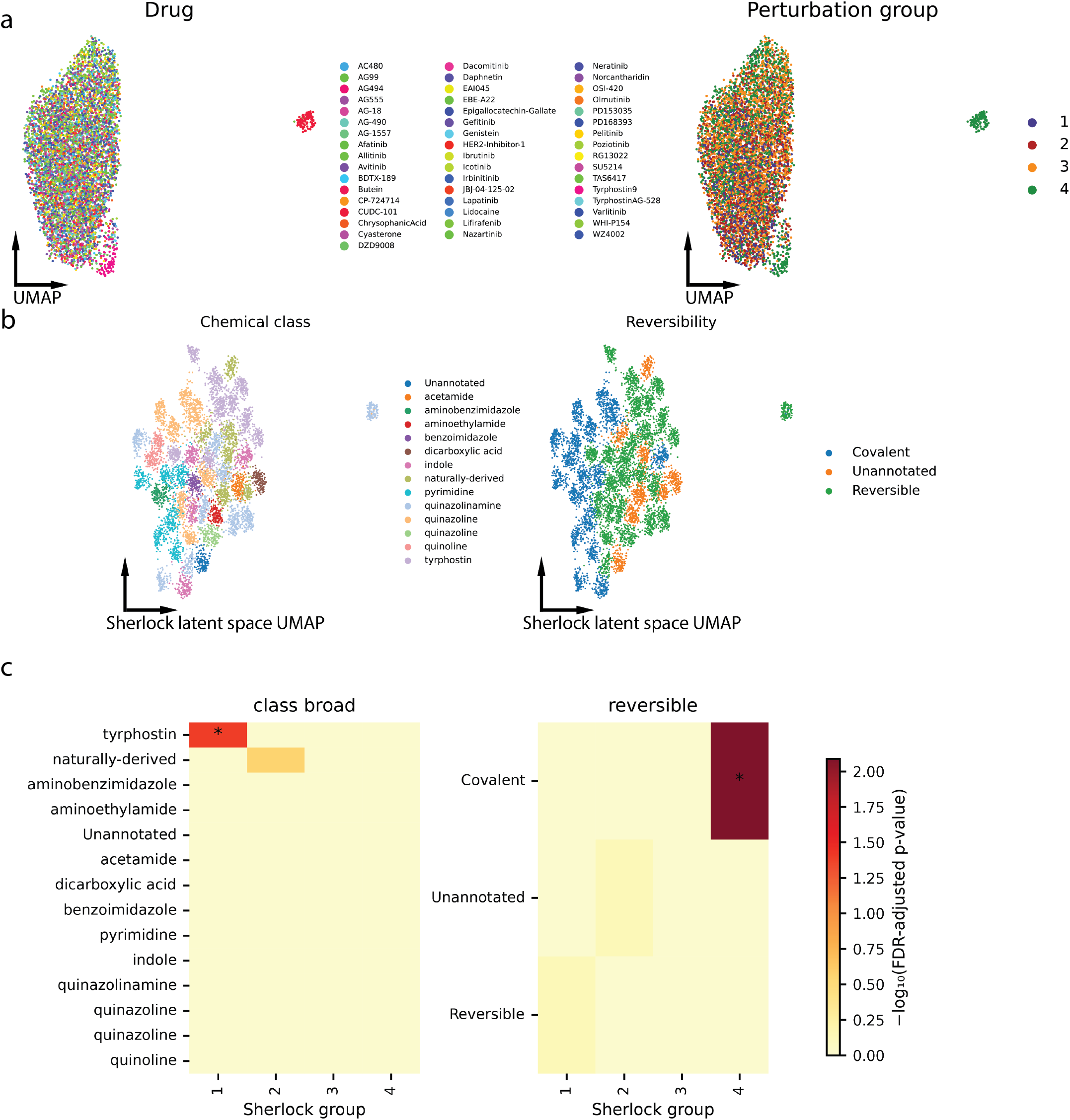
SHERLOCK perturbation groupings align with chemical and pharmacological drug properties. (a) UMAP embedding colored by drug names and perturbation group from Giglio et al. [24]. (b)UMAP of SHERLOCK latent embedding colored by chemical class and reversibility. (c) Enrichment of drug classes and reversibility categories across Sherlock perturbation groups. Heatmap colors indicate − log_10_(FDR-adjusted *p*-value), with asterisks marking significant enrichments at FDR *<* 0.05.

**Figure S3:**
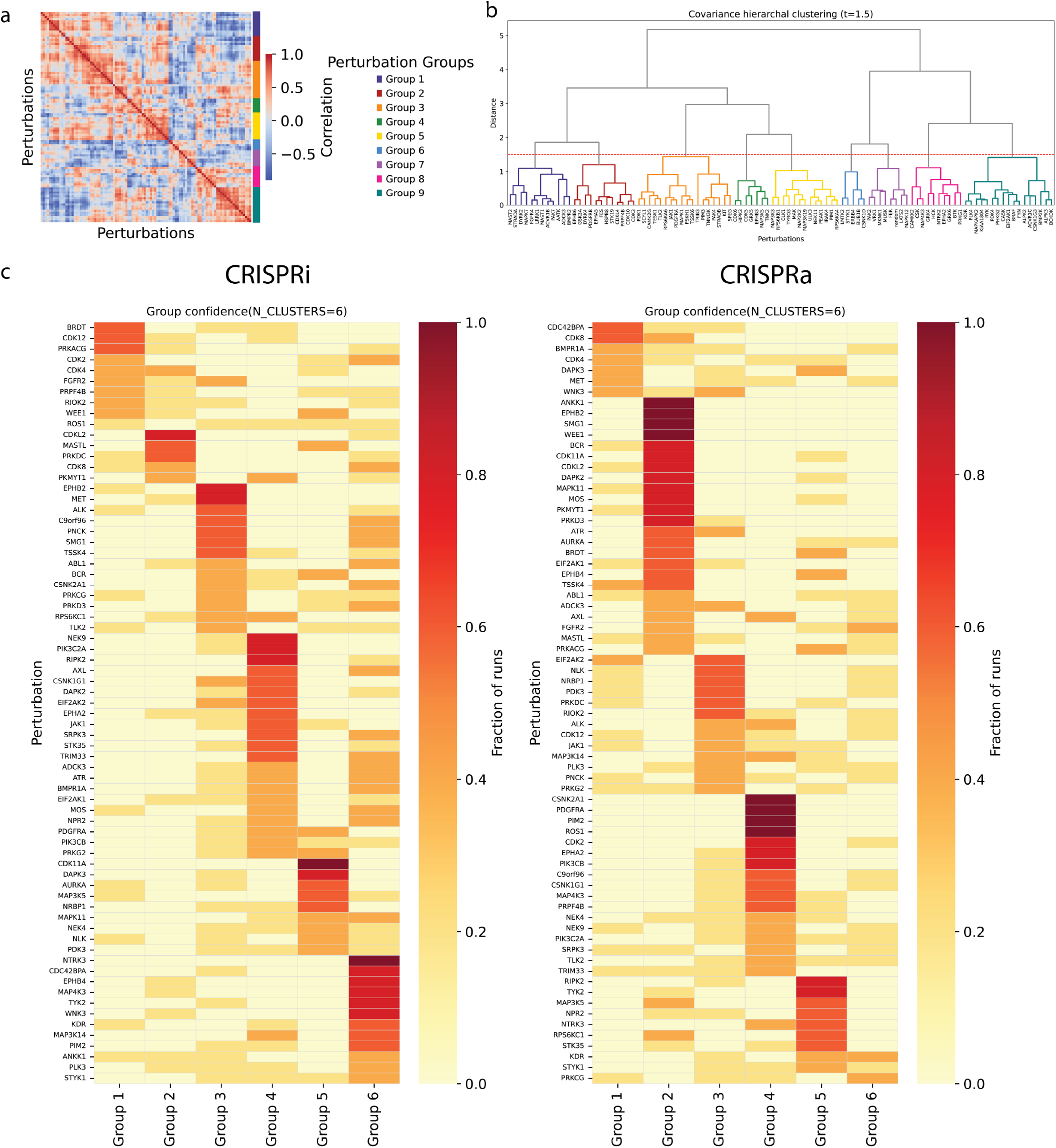
Robust and reproducible clustering of kinase perturbations across SHERLOCK runs. (a) Perturbation correlations derived from posterior row-covariance matrix (Σ_*p*_) of perturbation embeddings from the CRISPRa drug-target screen, revealing pathway-level block structure among genetic perturbations. (b) Hierarchical clustering of CRISPRa perturbation embeddings identifies coherent groups of functionally related genes, consistent with the covariance-based organization learned by SHERLOCK. c, Cluster assignment confidence across repeated SHERLOCK runs for CRISPR interference (CRISPRi; left) and CRISPR activation (CRISPRa; right). Each row represents a kinase perturbation, and each column represents one of the six consensus groups. Color intensity denotes the fraction of independent SHERLOCK runs in which a perturbation was assigned to each group, demonstrating reproducible clustering across both perturbation modalities.

**Figure S4:**
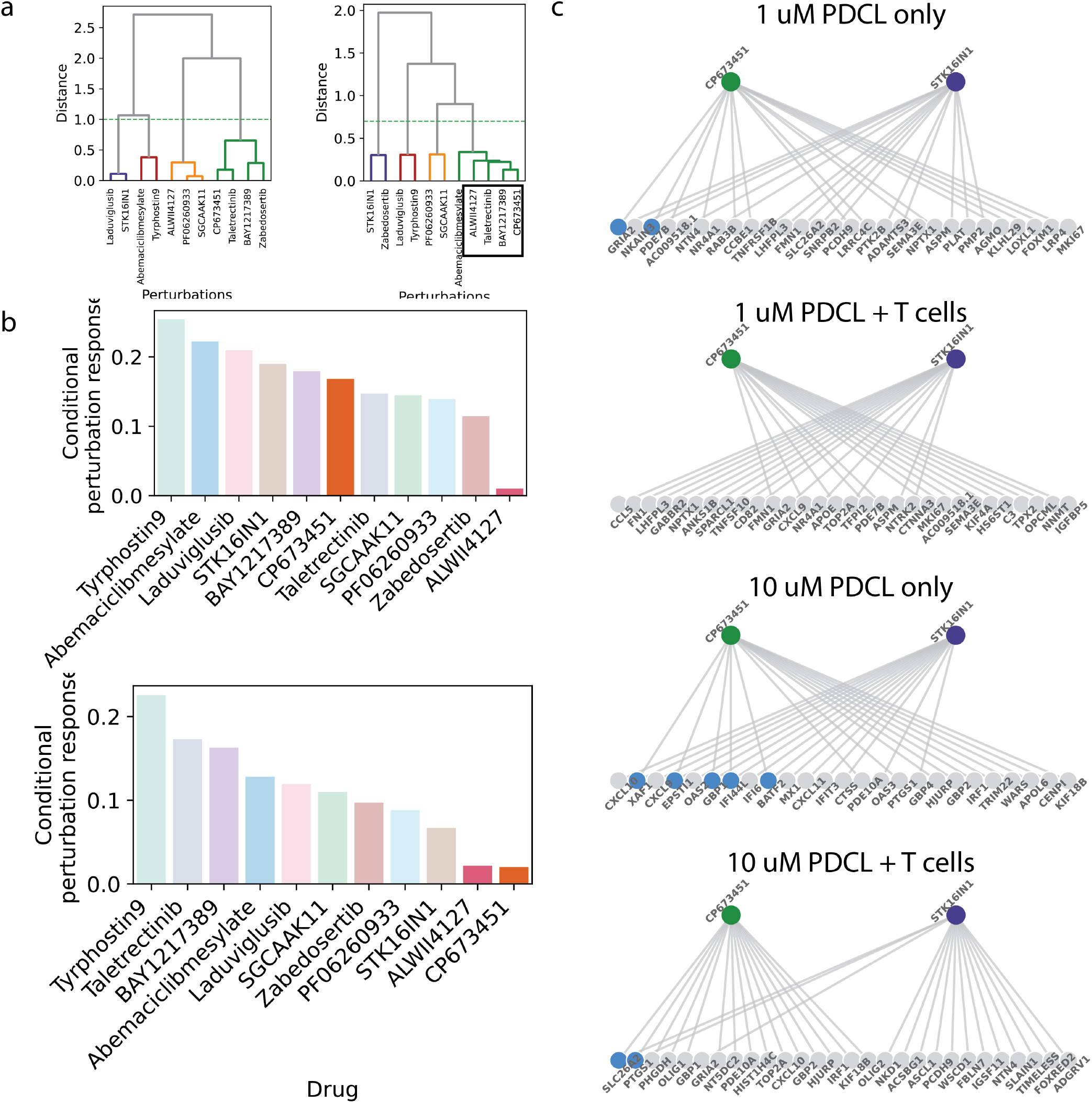
Condition-dependent perturbation effects and causal estimation of downstream gene-expression effects by SHERLOCK. (a) Hierarchical clustering of perturbations based on correlation derived from the learned covariance matrix under different conditions, revealing grouping structure that varies with T cell presence. Dendrograms illustrate perturbation-specific relationships between drugs (Left 1*uM*, Right 10*uM*). (b) Bar plot quantifying the condition-dependence of each drug’s perturbation effect (separation of drug-treated cells by T-cell condition). For each compound, the mean silhouette score across all pairwise T-cell conditions is computed in the model’s latent space, where a higher score indicates that the drug’s effect is strongly modulated by immunological context (i.e., cells cluster by condition within that perturbation), and a score near zero indicates a condition-invariant response. (c) Causal estimation of downstream genes for representative perturbations (CP673451 and STK16IN1) across conditions (No T cells and 0.5 T cell condition). Edges represent inferred perturbation–target relationships, and node color indicates shared or distinct targets.

**Figure S5:**
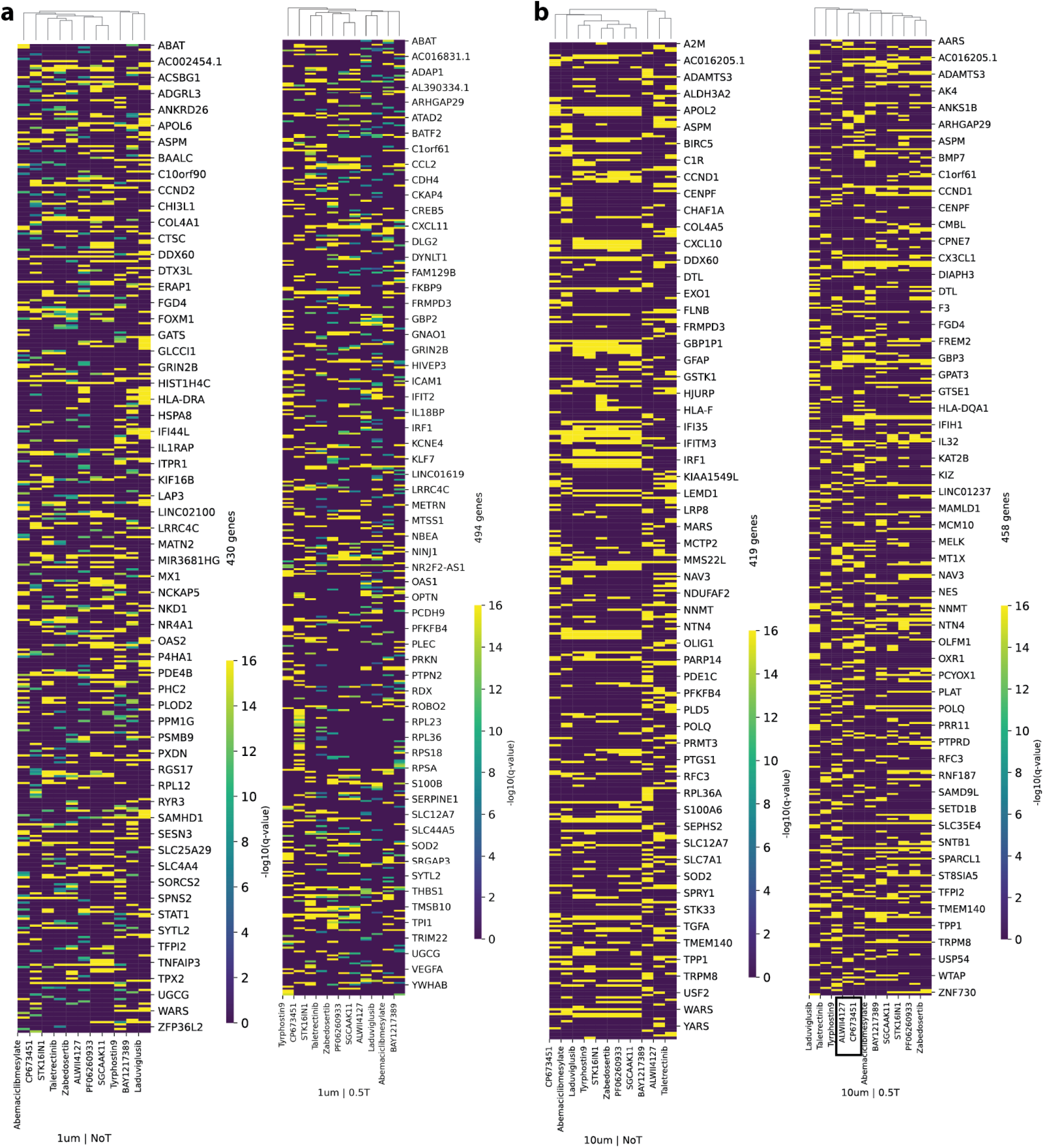
Causal estimation of downstream gene-expression effects across doses and T-cell co-culture conditions using SHERLOCK. (a) Heatmaps showing the statistical significance of causally estimated downstream gene-expression effects at low dose (1*µ*M) in the absence (NoT) and presence (0.5T) of T cells. Rows represent downstream response genes and columns represent drugs, hierarchically clustered based on SHERLOCK’s causally estimated expression effects. Color encodes log_10_(FDR-adjusted *q*-value) for each drug–gene pair. Only genes reaching *q <* 0.05 in at least one drug treatment are shown. (b) Corresponding heatmaps for high-dose (10*µ*M) conditions.

**Figure S6:**
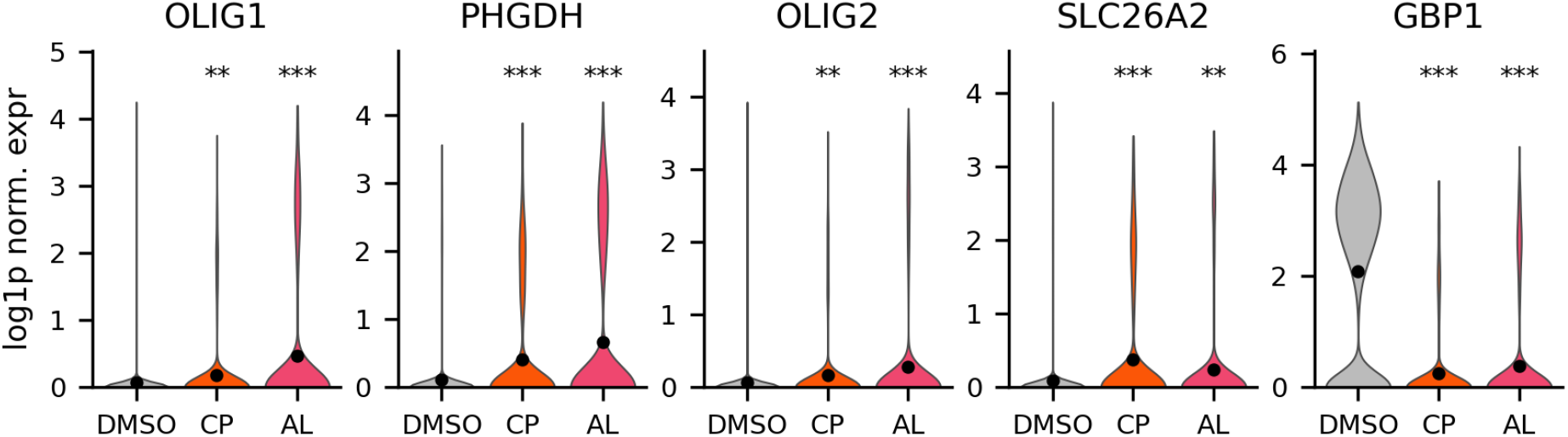
Expression of shared CP673451 and ALWII4127 targets in T-cell co-culture. Per-cell expression (log-normalized) of the shared target genes from the bipartite graph in BT333 cells co-cultured with T cells and treated with DMSO, CP673451 or ALWII4127 (10 *µ*M). Dots show means. Wilcoxon rank-sum test vs DMSO, Benjamini–Hochberg corrected: ^**^ *p <* 0.01, ^***^ *p <* 0.001. **DMSO** (*n* = 9,253 cells): reference. **CP673451** (*n* = 851 cells): OLIG1, *p* = 5.2 × 10^*−*3^; PHGDH, *p* = 1.9 × 10^*−*13^; OLIG2, *p* = 6.0 × 10^*−*3^; SLC26A2, *p* = 5.8 × 10^*−*12^; GBP1, *p* = 4.6 × 10^*−*183^. **ALWII4127** (*n* = 964 cells): OLIG1, *p* = 1.3 × 10^*−*11^; PHGDH, *p* = 5.6 × 10^*−*25^; OLIG2, *p* = 1.3 × 10^*−*4^; SLC26A2, *p* = 8.1 × 10^*−*3^; GBP1, *p* = 2.6 × 10^*−*174^.

**Figure S7:**
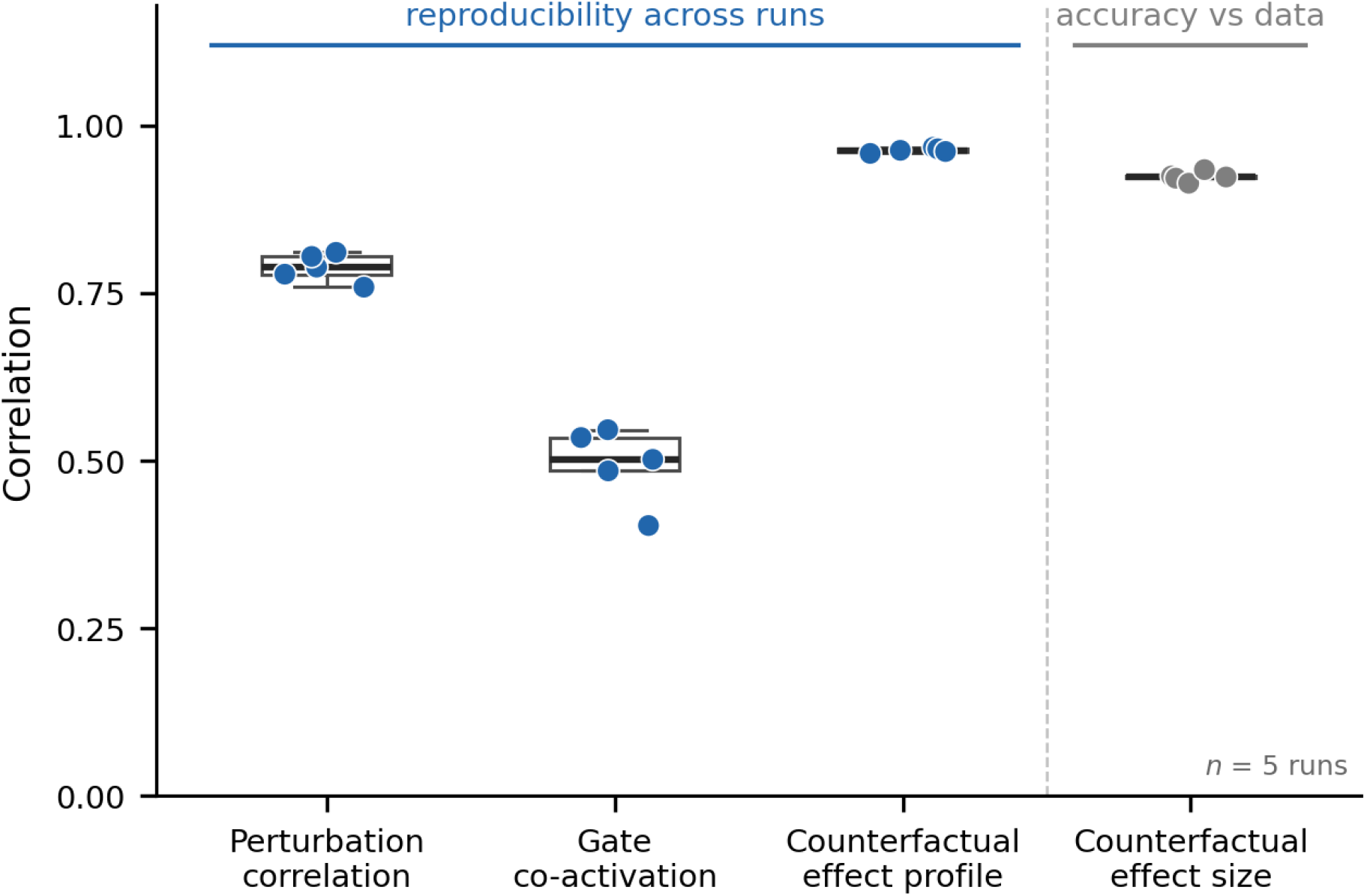
Reproducibility across five independently trained runs (Norman screen[43]). Each point is one run; the box shows the spread of the five values. The first three columns measure reproducibility across runs: each run is compared with the consensus of the other four (leave-one-out, Spearman). The fourth measures accuracy against the data within each run (Pearson). *Perturbation correlation*: the 105 × 105 matrix of learned similarities between perturbations (0.788 ± 0.021). *Gate co-activation*: for every pair of perturbations, how much their sets of gated latent factors overlap; this is compared instead of the gates themselves because latent coordinates are exchangeable between runs (0.494 ± 0.056). *Counterfactual effect profile*: the predicted perturbation-by-gene effect matrix (0.963 ± 0.004). *Counterfactual effect size*: the predicted per-perturbation effect magnitudes against the measured ones (0.924 ± 0.007).

**Figure S8:**
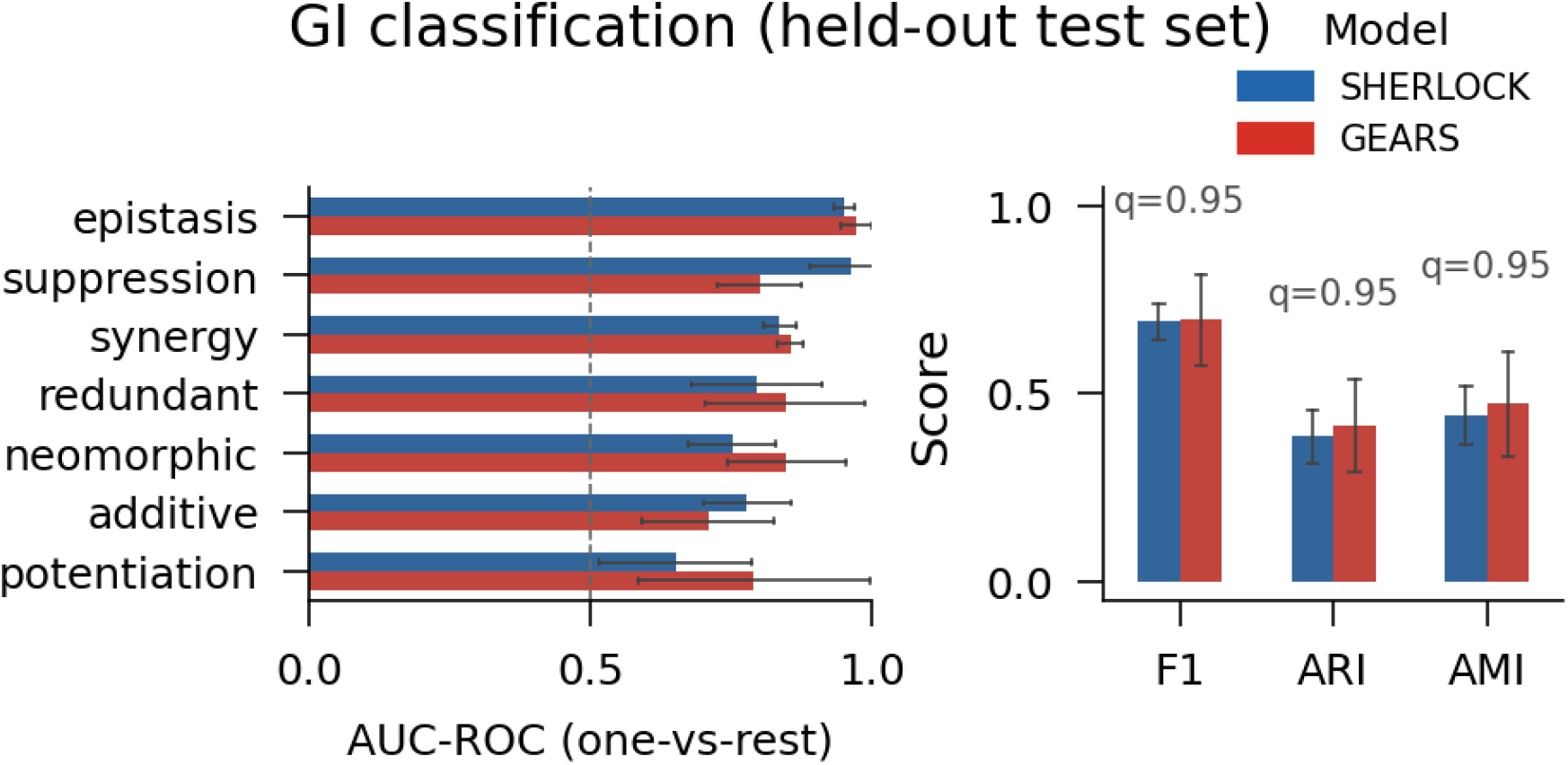
Genetic-interaction classification on the 22 held-out combinations. Each held-out pair’s effect is constructed from its two single perturbations, so neither model sees the pair’s own cells; the classes are then assigned by the same label-free routine used for the full panel, with the two synergy subtypes merged. Bars are mean ± SD over five independently trained runs. Left, per-class one-versus-rest AUC-ROC; right, weighted F1, ARI and AMI. No statistically significant difference in weighted F1 was detected between SHERLOCK and GEARS (weighted F1 0.692 0.047 vs 0.696 0.120, *q* = 0.95), and with 22 pairs spread over seven classes; the per-class supports are in the low single digits, limiting the precision of class-specific performance estimates.

**Figure S9:**
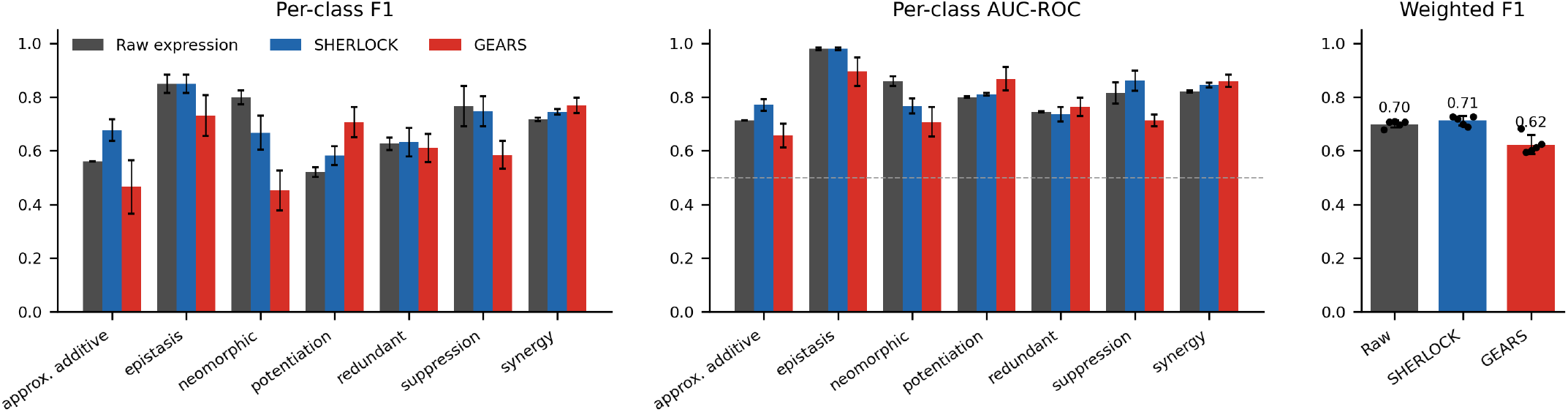
Genetic-interaction classification from measured expression, SHERLOCK and GEARS. All three effect sources were scored with the same Theil-Sen regression and label-free classifier on the 86 labeled pairs, with the synergy subtypes merged. Measured expression involves no training, so its five runs differ only in the regression seed, whereas SHERLOCK and GEARS use their five trained runs. Bars are mean ± SD over runs. Left, per-class F1; middle, per-class one-versus-rest AUC-ROC (dashed line, chance); right, weighted F1 with individual runs as points.

**Figure S10:**
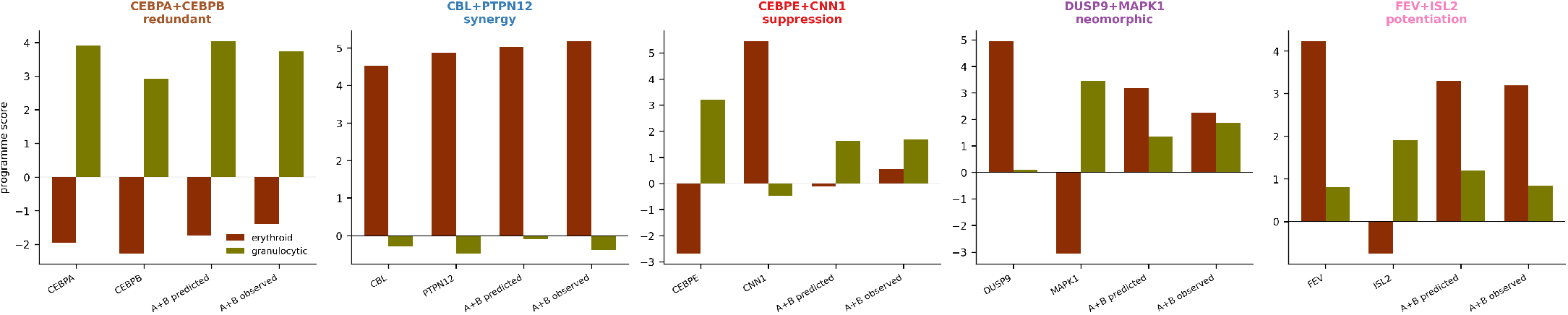
Worked examples of five labeled pairs. Measured erythroid and granulocytic program scores for each parent, the model’s predicted combination and the measured combination. A score is the mean change of that program’s genes relative to control, in units of the profile’s own gene-wise spread, so positive means the program is switched on and negative that it is pushed below control. All five pairs were in the model’s training set, so this shows fit rather than prediction.

**Figure S11:**
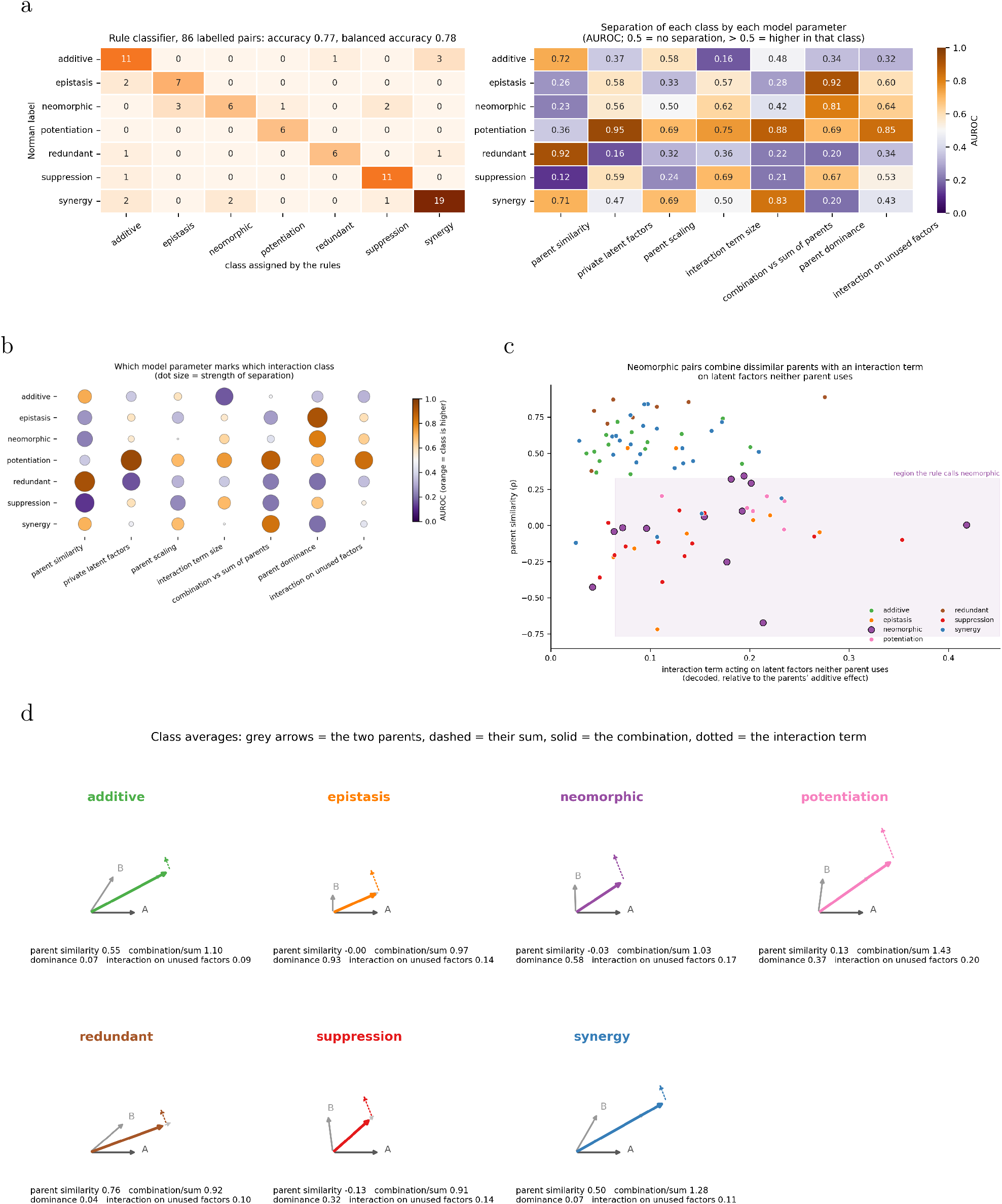
Interaction class read from the model’s parameters. (a) Confusion matrix of the seven-rule decision list on the 86 labeled pairs (accuracy 0.77, balanced 0.78; rows, reference labels; columns, assigned class), and how well each parameter separates each class on its own (one-versus-rest AUROC). Cut points were fit on these pairs, so both panels describe separation rather than predictive performance. (b) The same AUROCs as a fingerprint per class: color gives direction (orange, higher in that class; purple, lower), dot size the strength of separation. (c) The neomorphic rule: each pair placed by the decoded effect of its interaction term on latent factors neither parent uses and by parent similarity *ρ*, with the rule’s region shaded. (d) Schematic illustrations of the seven classes drawn from their mean parameters: grey arrows are the single perturbations, separated by arccos of their mean similarity and scaled by mean dominance; dashed, their sum; solid, the combination; dotted, the interaction on unused factors (its direction is arbitrary).

**Figure S12:**
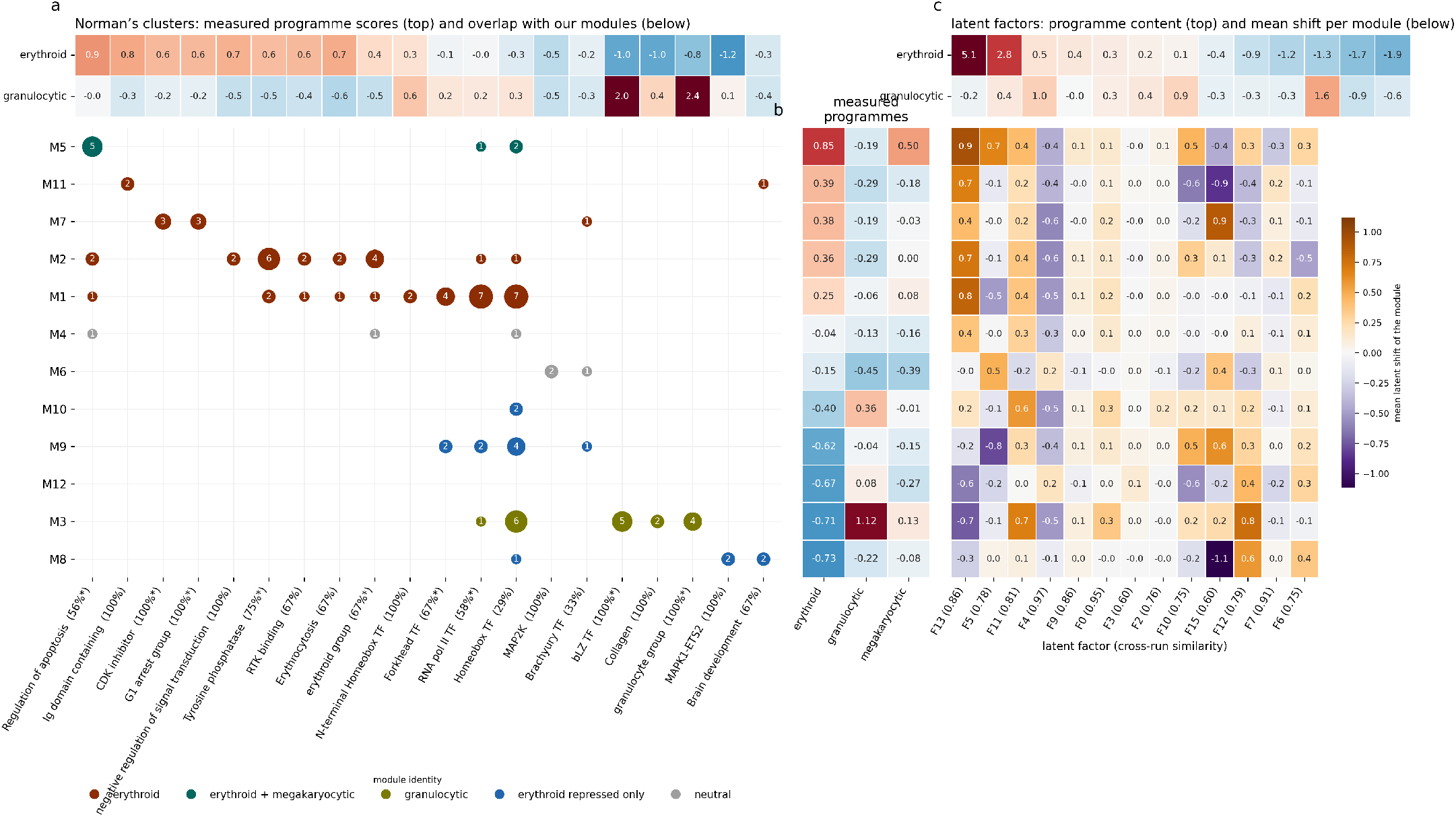
Consensus modules against Norman’s clusters, measured programs and latent factors. (a) Overlap between the consensus modules (rows) and Norman et al.’s perturbation-map clusters and phenotype groups (columns); dot size and number give the members shared, dot color the module’s identity, and the percentage under each cluster the share of its members in its best module (asterisk, BH *q <* 0.05 against random modules of the same sizes). The strip above gives each cluster’s own measured erythroid and granulocytic scores. (b) Mean measured erythroid, granulocytic and megakaryocytic scores per module. The megakaryocytic score is a weak signal: scoring independent halves of each perturbation’s cells agrees at 0.84, against 0.99 and 0.97 for the other two lineages. (c) Mean shift of each module along each reproducible latent factor, with the strip above giving each factor’s erythroid and granulocytic content; factors are labeled with their cross-run signature similarity in parentheses.

**Figure S13:**
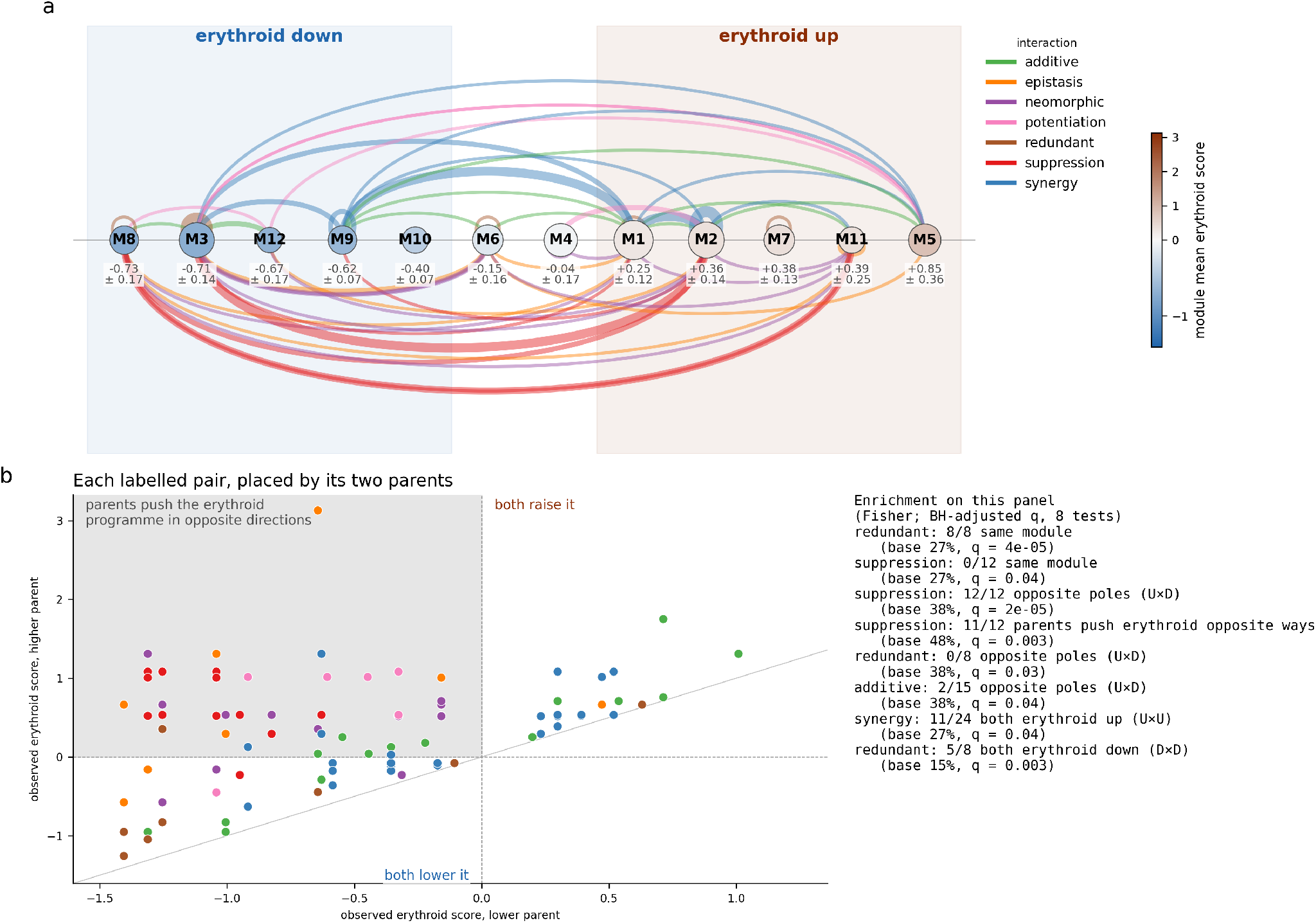
labeled interactions placed on the erythroid axis. (a) Each labeled pair is drawn as an arc between the consensus modules of its two parents. Modules run from erythroid down on the left to erythroid up on the right, with their mean observed erythroid score and its standard error over member perturbations printed below each one; node color repeats that score (color bar) and node size scales with the number of members. Arc color gives the interaction class: approximately additive, synergistic, redundant and potentiating pairs are drawn above the line, suppressive, epistatic and neomorphic pairs below, and arcs are semi-transparent so that overlapping pairs remain visible. Of the modules, M3 is granulocytic and M5 is erythroid plus megakaryocytic; the rest are erythroid-high, neutral or erythroid-low as described in the text. (b) The same pairs placed by the observed erythroid scores of their two parents (lower parent on the *x*-axis, higher on the *y*-axis); the inset lists the BH-adjusted Fisher tests reported in the text.

**Figure S14:**
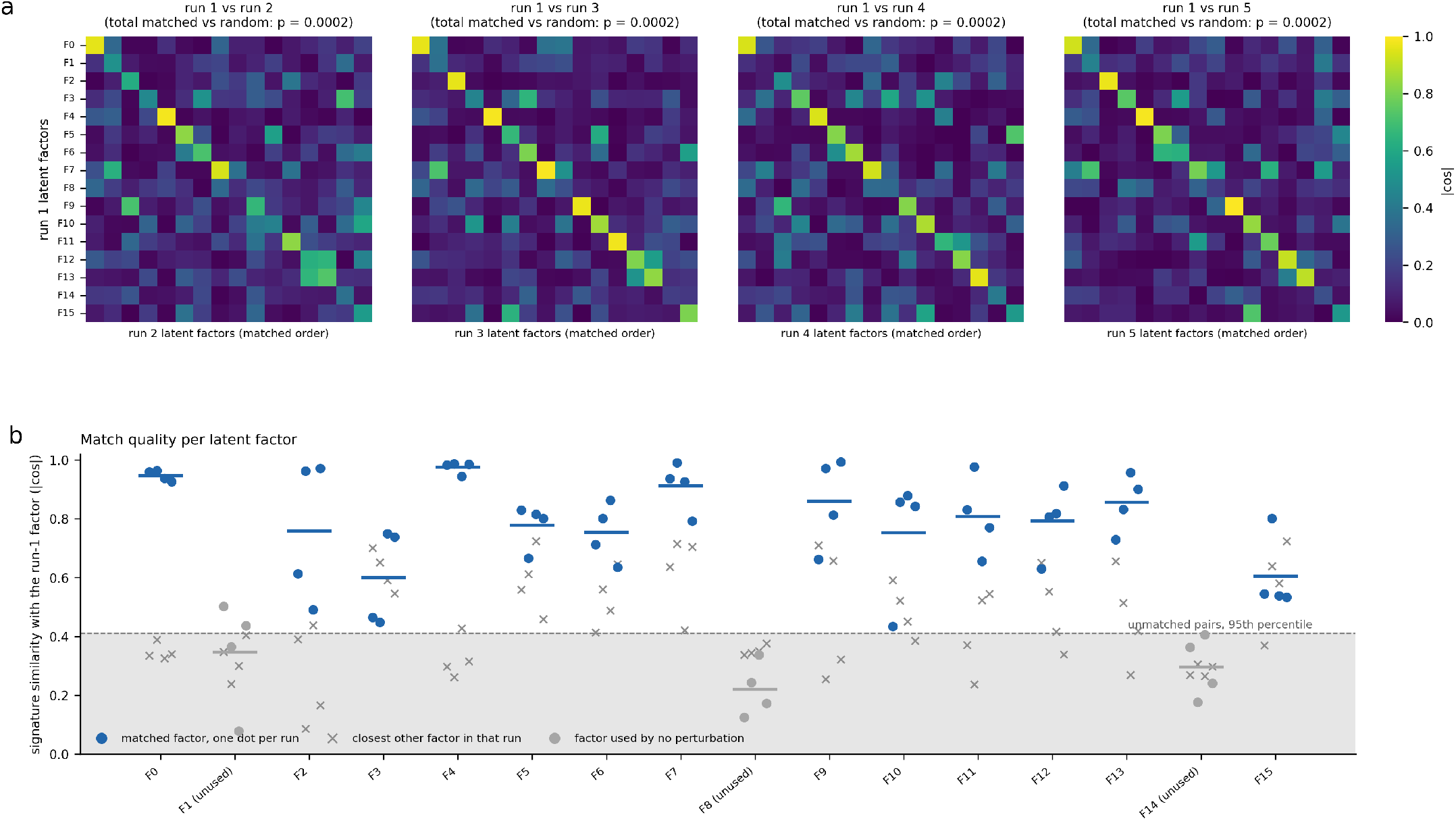
Latent factors match across independently trained runs. Factors are identified by their decoded gene signature, the expression change produced by moving along the factor. (a) Absolute cosine similarity between the signatures of runs 2–5 and those of run 1, with columns reordered by the Hungarian matching; a clean diagonal means each factor has one clear counterpart. Each run’s total matched similarity exceeds 5,000 random pairings (*q* = 0.0002). (b) Per factor, the similarity to its matched partner in each run (dots), the closest other factor in that run (grey crosses) and the range spanned by unmatched pairs (shaded; 95th percentile marked). Factors used by no perturbation are grey; F3 and F15, whose closest alternative is as similar as their match, are not interpreted.

**Figure S15:**
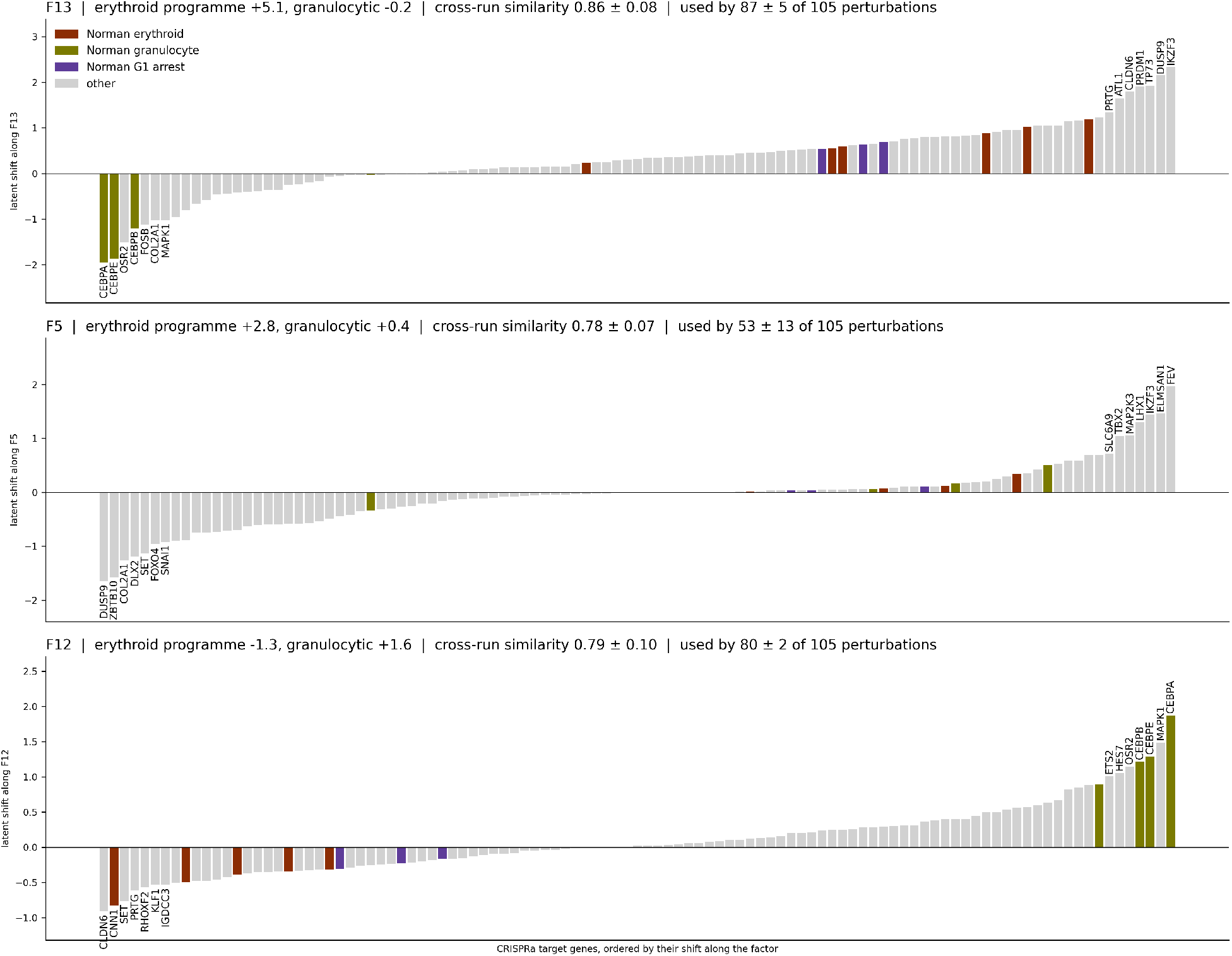
Which perturbations drive the three interpretable latent factors. Shift of each CRISPRa target along F13 (erythroid), F5 (erythroid) and F12 (granulocytic versus erythroid), one bar per target gene, ordered by that shift and colored by Norman phenotype group; the most extreme genes on each side are named. Panel titles give the factor’s program content, its cross-run signature similarity and the number of perturbations using it, each ± SD over the five runs.

